# UNC-5 (UNC5) mediates neuronal outgrowth patterning in *Caenorhabditis elegans* by regulating UNC-40 (DCC) asymmetric localization

**DOI:** 10.1101/083436

**Authors:** Gerard Limerick, Xia Tang, Won Suk Lee, Ahmed Mohamed, Aseel Al-Aamiri, William G. Wadsworth

## Abstract

Neurons extend processes that vary in number, length, and direction of outgrowth. Extracellular cues help determine outgrowth patterns. In *Caenorhabditis elegans,* neurons respond to the extracellular UNC-6 (netrin) cue via UNC-40 (DCC) and UNC-5 (UNC5) receptors. Previously we presented evidence that UNC-40 asymmetric localization at the plasma membrane is self-organizing and that UNC-40 can localize and mediate outgrowth at randomly selected sites. We also postulate that the process is statistically dependent, *i.e.* if the probability of outgrowth at one site changes then the probability at another site(s) must also change. Over time, the direction of outgrowth activity fluctuates across the membrane. A probability distribution describes the likelihood of outgrowth in each direction. Random walk modeling predicts that the degree to which the direction of outgrowth fluctuations affects the outward displacement of the membrane. We predict that extracellular cues create patterns of outgrowth by differentially affecting the degree to which the direction of outgrowth activity fluctuates along the membrane. This produces different rates of outgrowth along the surface and creates patterns of extension. Here we present evidence that UNC-5 (UNC5) receptor activity regulates UNC-40 asymmetric localization and the patterning of outgrowth. We show that *unc-5* mutations alter UNC-40 asymmetric localization and the patterns of outgrowth that neurons develop. Genetic interactions suggest UNC-5 acts through the UNC-53 (NAV2) cytoplasmic protein to regulate UNC-40 asymmetric localization in response to both the UNC-6 and EGL-20 (wnt) extracellular cues.

## Introduction

During development, an intricate network of neuronal connections is established. As processes extend from the neuronal cell bodies, distinct extension patterns emerge. Some extensions remain as a single process, whereas others branch and form multiple processes. If they branch, the extensions can travel in the same or in different directions. Extensions vary in length. Extracellular cues are known to influence this patterning, but the underlying logic that governs the formation of patterns remains a mystery.

The secreted extracellular UNC-6 (netrin) molecule and its receptors, UNC-5 (UNC5) and UNC-40 (DCC) are highly conserved in invertebrates and vertebrates, and are known to play key roles in cell and axon migrations. In *Caenorhabditis elegans*, UNC-6 is produced by ventral cells in the midbody and by glia cells at the nerve ring in the head (WADSWORTH *et al.* 1996; WADSWORTH AND HEDGECOCK 1996; ASAKURA *et al.* 2007). It’s been observed that neurons that express the receptor UNC-40 (DCC) extend axons ventrally, towards the UNC-6 sources; whereas neurons that express the receptor UNC-5 (UNC5) alone or in combination with UNC-40 extend axons dorsally, away from the UNC-6 sources (HEDGECOCK *et al.* 1990; LEUNG-HAGESTEIJN *et al.* 1992; CHAN *et al.* 1996; WADSWORTH *et al.* 1996).

It is commonly proposed that axons are guided by attractive and repulsive mechanisms (Tessier-Lavigne and Goodman 1996). According to this model, an extracellular cue acts as an attractant or repellant to direct neuronal outgrowth towards or away from the source of a cue. UNC-5 (UNC5) has been described as a “repulsive” netrin receptor because it mediates guidance away from netrin sources (Leung-Hagesteijn et al. 1992; Hong et al. 1999; Keleman and Dickson 2001; Moore et al. 2007). The attraction and repulsion model is deterministic. That is, given the same conditions, the response of the neuron, attractive or repulsive, will always be the same. This idea forms the bases of the analysis and interpretation of experimental results. Axonal growth cone movement towards or away from the source of a cue is considered to be mediated by attractive or repulsive responses to the cue. In genetic studies, a mutation that disrupts movement towards the cue source denotes gene function within an attractive pathway, whereas mutations that disrupt movement away from a source denotes gene function within a repulsive pathway. If an axonal growth cone is observed to move towards and then away from the source of a cue, the responsiveness of a neuron is thought to switch from attractive to repulsive. However, it is important to note that attraction or repulsion is not an intrinsic property of the interaction between the receptor and ligand. In fact, the interaction only promotes or inhibits outward movement of the membrane. Attraction and repulsion refers to a direction, which is an extrinsic property of the cellular response that varies depending on the physical positions of the ligands. Movement towards or away from a cue source is caused by attractive and repulsive *effects*, such as chemoattraction and chemorepulsion, which is movement that is directed by chemical gradients of ligands. We argue that classifying gene function as attractive or repulsive is problematic since attraction and repulsion are not intrinsic properties of cellular mechanisms.

We have proposed an alternative model in which the movement of neuronal outgrowth is not considered in terms of attraction and repulsion. This model comprises three concepts. The first concept is that receptors along the surface of the membrane change position. This is important since the spatial distribution of receptors can influence the movement that a neuron has in response to the extracellular ligands (NGUYEN *et al.* 2014; NGUYEN *et al.* 2015). We hypothesize that the spatial distribution of UNC-40 can influence the manner though which force is applied to the membrane and thereby affect the outward movement of the membrane. It’s known that the surface localization of the UNC-40 receptor undergoes dramatic changes during the development of the HSN axon (ADLER *et al.* 2006; XU *et al.* 2009; KULKARNI *et al.* 2013). As HSN axon formation begins, UNC-40 becomes asymmetrically localized to the ventral surface of the cell body, which is nearest the ventral sources of the secreted UNC-6 ligand. Live imaging of the developing leading edge reveals a dynamic pattern of UNC-40 localization with areas of concentrated UNC-40 localization shifting positions along the surface (KULKARNI *et al.* 2013). Dynamic UNC-40::GFP localization patterns have also been reported during anchor cell extension (WANG *et al.* 2014). Similar to axon extension, the anchor cell also sends an extension through the extracellular matrix and this extension is also regulated by UNC-40 and UNC-6 (ZIEL *et al.* 2009; HAGEDORN *et al.* 2013). Live imaging of the anchor cell reveals that UNC 40::GFP “clusters” form, disassemble, and reform along the anchor cell’s plasma membrane (WANG *et al.* 2014).

The second concept is that the asymmetric localization of the receptor, and the subsequent outgrowth activity it mediates, is stochastically oriented. It was observed that UNC-40 can asymmetrically localize to a randomly selected surface if the UNC-6 ligand is not present to provide a pre-established asymmetric cue (XU *et al.* 2009). We noted that the self-organizing nature of UNC-40 localization is reminiscent of a self-organizing process observed in single-cell yeast, *Dictyostelium discoideum*, and neutrophils, where cell movement will occur in a random direction if the chemotactic cue is absent or is uniformly presented (FRASER *et al.* 2000; ARRIEUMERLOU AND MEYER 2005; MORTIMER *et al.* 2008). The process through which outgrowth activity becomes asymmetrically organized is thought to utilize positive- and negative-feedback loops (BOURNE AND WEINER 2002; GRAZIANO AND WEINER 2014). Such loops might also drive the asymmetric localization of UNC-40 (XU *et al.* 2009; WANG *et al.* 2014). Positive and negative feedback are considered complementary mechanisms; positive feedback amplifies the polarized response to an extracellular cue, while negative feedback limits the response and can confine the positive feedback to the leading edge (BOURNE AND WEINER 2002). The biological nature of feedback loops controlling UNC-40 activity is unclear. However, they may involve the differential transport of receptors and effectors to the plasma membrane surface. Imaging experiments of cells in culture suggest that netrin-1 (UNC-6) regulates the distribution of DCC (UNC-40) and UNC5B (UNC-5) at the plasma membrane (GOPAL *et al.* 2016). In these studies, netrin-1 (UNC-6) was shown to stimulate translocation of DCC (UNC-40) and UNC5B (UNC-5) receptors from intracellular vesicles to the plasma membrane and, further, the transported receptors were shown to localize at the plasma membrane (GOPAL *et al.* 2016).

We argue that the process that localizes UNC-40 to a site on the plasma membrane possesses inherent randomness (Figure 1). Evidence suggests that the conformation of the UNC-40 molecule controls whether the process will cause UNC-40 localization to the site of UNC-6 interaction or to another site (XU *et al.* 2009). We observed that a single amino acid substitution in UNC-40 will allow UNC-40 to asymmetrically localize to different surfaces in the absence of UNC-6. The binding of UNC-6 to this UNC-40 molecule causes localization to the surface nearest the UNC-6 source. However, the binding of UNC-6 with a single amino acid substitution will enhance the asymmetric localization to different surfaces. A second-site UNC 6 amino acid substitution will suppress this enhancement and increase UNC-40 asymmetric localization at the surface towards the UNC-6 source. These results indicate that UNC-40 conformational changes differentially influence each activity. In the context of feedback loops, UNC-40 activity regulates both the positive and negative loops that control the asymmetric localization of UNC-40 to the plasma membrane. Because the system is controlled by the conformation of the molecule, randomness will be introduced in the system by stochastic fluctuations in ligand-receptor binding and by stochastic conformational changes.

**Figure 1.**
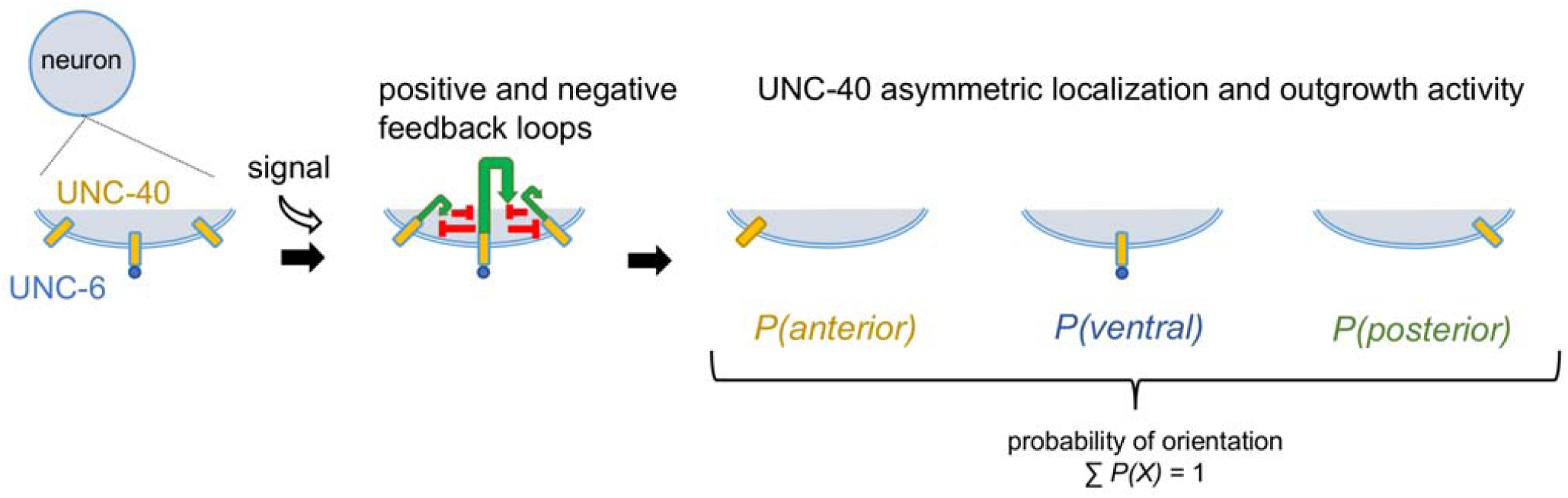
Statistically dependent asymmetric localization (SDAL). At sites along the plasma membrane, UNC-40 interacts with the UNC-6 extracellular cue. A self-organizing process is triggered that utilizes positive- and negative-feedback loops. Positive feedback (green arrows) amplifies the polarized response to an extracellular cue, while negative feedback (red lines) limits the response and can confine the positive feedback to the site of UNC-40 and UNC-6 interaction. The outcome of an UNC-40 receptor’s activity is to either cause an UNC-40 receptor to localize and mediate outgrowth at the site of UNC-6 interaction or at a different site. Randomness is considered inherent in this process and each localization event is mutually exclusive. This statistical dependence means that the probability of UNC-4O localizing and mediating outgrowth at the site of UNC-6 interaction affects the probability of UNC-40 localizing and mediating outgrowth at another site, and vice versa. As time passes, this process causes randomly directed outgrowth activity (force) that drives the outward movement of the membrane.

The outcome of an UNC-40 receptor’s activity is to either cause an UNC-40 receptor to localize to the site of UNC-6 interaction or to a different site (Figure 1). This is an important discovery because it means that the asymmetric localization events are mutually exclusive and, therefore, there is statistical dependence. We refer to this process as ‘statistically dependent asymmetric localization’ (SDAL). This model states that the probability of UNC-4O localizing and mediating outgrowth at the site of UNC-6 interaction affects the probability of UNC-40 localizing and mediating outgrowth at another site, and vice versa. We have found that other extracellular cues can also affect UNC-40 asymmetric localization, and thus can influence the probability of UNC-40-mediated outgrowth from different sites (TANG AND WADSWORTH 2014; YANG *et al.* 2014).

The development of an extension can be considered as a stochastic process. At any one instance of time at innumerable sites along the neuron’s surface, UNC-40 interacts with UNC-6 to mediate UNC-40 asymmetric localization and the outgrowth response. At the next instance of time, other UNC-40 receptors, including any just transported to the surface, may interact with UNC-6. Because the plasma membrane is a fluid, the forces generated by the outgrowth response are not always acting in parallel and the direction of outward force can fluctuate (Figure 2A). It is the collective impact of all outgrowth events over a period of time that allows the extension to form. The outward movement of an extension could be precisely described if the effect of each outgrowth event where known. However, it is extremely difficult to measure the effect of each single event since there are innumerable events happening at each instance of time. We also argue that the pattern of UNC-40 localization and outgrowth across the surface of the membrane evolves over time through a random process. Therefore, the outgrowth events can only be described probabilistically, and as such the time evolution of extension is also probabilistic in nature.

**Figure 2.**
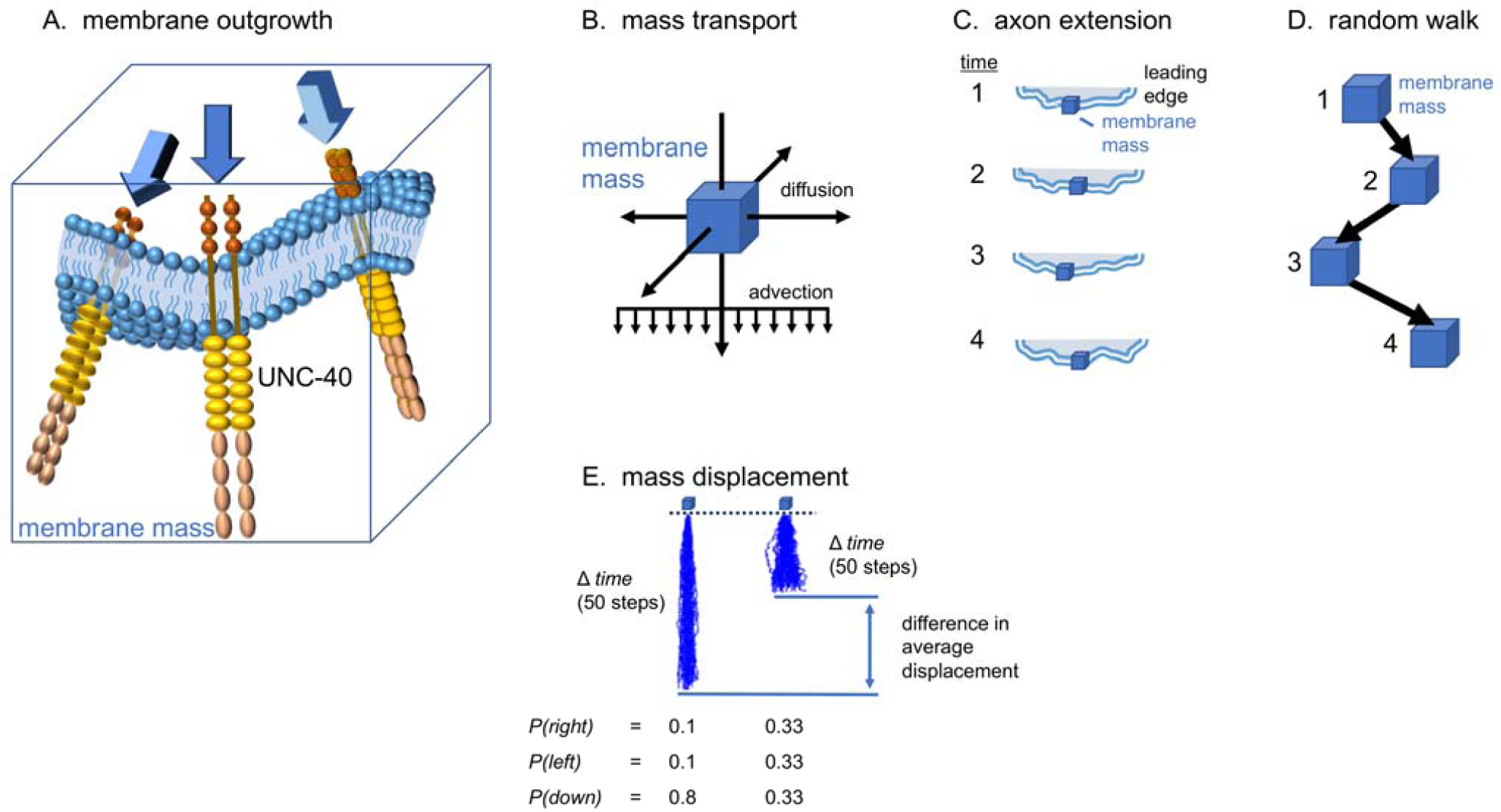
Model for outgrowth movement. **(A)** The outward movement of the neuronal membrane is depicted as a mass transport phenomena. The cell membrane is fluid and membrane mass will move in different directions as the membrane is subjected to forces (arrows) which change its shape. Receptors mediate cellular responses which creates the outward force. The force causes movement of the lipids and proteins of the plasma membrane. A unit of this mass is shown within a box. Membrane mass is represented by a box in subsequent schematic diagrams. **(B)** The mean flow of membrane mass (box) can be described as advection and diffusion. The probability density function of the position of mass as a function of space and time is described mathematically by an advection-diffusion equation. Mass transport by a mean velocity field is advection. Because of the SDAL process and the fluid nature of the membrane, mass transport also occurs through random movement, which is diffusion. **(C)** During outward movement of the leading edge (times 1-4), membrane molecules move in the direction of advection as well as randomly in other directions. **(D)** The path that the membrane molecules take during outgrowth can be described as a random walk, which is a succession of randomly directed steps. Depicted are the position of mass after each step of a succession of four steps as shown in C. Each step corresponds to a time point. **(E)** For two examples, 50 simulated random walks of 500 steps were plotted from an origin (0,0). For each step the probability of moving to the right, left, or down is given below the plots. The plots illustrate that increasing the degree to which the direction of movement fluctuates, decreases the outward distance that mass can travel. We predict that the SDAL process influences the degree of random membrane movement and, consequently, the outward displacement of the membrane.

Understanding the role a gene plays in controlling outgrowth movement might require knowledge of its role in regulating this stochastic process. To do this, we use the direction of HSN extension. We reason that the collective impact of all the outgrowth events over a period of time cause the development of the HSN axon. The direction of extension from the cell body has a probability of being orientated in one direction (KULKARNI *et al.* 2013; TANG AND WADSWORTH 2014; X AND WG 2014; YANG *et al.* 2014). Mathematically, the direction of HSN extension is a variable that takes on different values; “anterior”, “posterior”, “ventral”, and “dorsal”. A probability is associated with each outcome, thus creating a probability distribution. This distribution describes the effect that all the outgrowth events had over a period of time. During normal development, the probability of each UNC-40-mediated outgrowth event being ventrally oriented is very high and a ventral extension develops. We have shown that certain gene mutations affect the probability distribution, thus revealing that a gene plays a role in the stochastic process. We can compare wildtype and mutants to gage the degree to which a mutation causes the direction of extension to fluctuate. This reflects the degree to which the mutation has caused the direction of outgrowth activity, and the outward force it creates, to fluctuate over the course of extension development.

We argue that understanding the function of a gene in terms of a stochastic model of membrane movement is useful. Often the goal of a genetic analysis of axon guidance is to uncover a molecular mechanism. Frequently, a deterministic model is made which describe some molecular event that the gene affects. Because the mutation affects axon guidance, the molecular event plays a role in causing directed movement. However, these models tend to reduce a complex biological process to an isolated component. In reality, understanding how a molecular event is able to cause directed movement requires knowledge of all the many ways in which the event influences, and is influenced by, the other molecular events of directed movement. A stochastic model is a useful tool of exploring how a gene affects the overall behavior of the system. To make an analogy, a roulette wheel can be described deterministically; if every force acting on the ball at every instance of time is known than the number on which the ball stops can be precisely determined. The role of a component of the roulette wheel could be described by the effect that it has on the forces which act on the ball at every instance of time. However, understanding how the effect of this component causes a particular outcome require understanding the effects of all the other components. Because this is so complex, the outcome of a roulette wheel is studied using a stochastic model. That is, how does the component affect the probability of the ball stopping on a particular number. A roulette wheel must be exactly levelled to have an equal probability for each number. Removing a component of the wheel can cause the wheel to tilt in a particular manner. This will result in a new outcome, *i.e.* the ball will have a higher probability of stopping on certain numbers. Although this does not reveal the precise event that occurs between the component and the ball, it will reveal the effect that the component has in determining an outcome. Further, by studying the effect of removing multiple components, relationships that lead to particular outcomes can be revealed.

The third concept of our model is that neuronal membrane outgrowth is a mass transport phenomena which can be described as advection and diffusion (Figure 2). Signaling by UNC-40 receptors along a surface of the neuron can lead to cytoskeletal changes which create force and membrane movement (Figure 2A). As a result, there is a mean flow of membrane mass in outward direction (Figure 2B). This motion is advection, which is mass transport by a mean velocity field. In addition to advection, membrane mass transport also occurs through random movement, *i.e.* diffusion. Because the cell membrane is fluid, membrane mass will move in different directions as the membrane is subjected to forces which change its shape (Figure 2C). The degree to which the membrane mass undergoes random movement is important because diffusion processes and advection processes have different effects on the extent to which mass will be displaced outward in a given amount of time. The random movements can be mathematically described using random walks. A random walk is a succession of randomly directed steps (Figure 2D). Random walk models are used to describe many diverse types of behavior, including the movement of a particle through fluid, the search pattern of a forging animal, and the fluctuating price of a stock. The behavior of neuronal growth cone movement during chemotaxis has also been modeled using random walks (KATZ *et al.* 1984; BUETTNER *et al.* 1994; ODDE AND BUETTNER 1995; WANG *et al.* 2003; MASKERY *et al.* 2004). However, rather than using a random walk model to describe the gross morphological changes observed during growth cone movement, in this study the random walk is used to model the random movement of membrane mass in order to understand how gene activity influences the outward displacement of the membrane. A property of random motion is that the mean square displacement grows proportionate to the time traveled. This means that the more the direction of movement fluctuates, the shorter the distance of travel in a given amount of time (Figure 2E). The model predicts that if force is applied to the membrane in a manner that increases random movement then the outward displacement of the membrane’s mass will decrease.

Because of the effect random movement has on displacement, the SDAL model makes predictions about how UNC-40 activity affects the rate of extension. In a deterministic model, outgrowth activity causes straight-line outward motion from the site of interaction. The SDAL model predicts that the interaction between UNC-40 and UNC-6 increases the probability that UNC-40 asymmetric localization and UNC-40-mediated outgrowth will be oriented at the site of interaction. It also decreases the probability that localization and outgrowth will be oriented elsewhere. Therefore, the interaction influences the spatial distribution of UNC-40 along the surface and, in doing so, will change the way forces are applied to the membrane. As this process continues over time, the direction of the forces acting on the fluid membrane fluctuates. This will alter the advective and diffusive transport of membrane mass. As an example, if the probability of ventral outgrowth is 0.33, of anterior outgrowth is 0.33, and posterior outgrowth is 0.33 then there will be a high degree of random movement. Interactions between UNC-40 and UNC-6 at the leading ventral surface could shift the probabilities for ventral outgrowth to 0.8, for anterior outgrowth to 0.1, and for posterior outgrowth to 0.1. This change will decrease the degree to which the direction of outgrowth fluctuates. As modeled in Figure 2E, this will increase displacement, meaning that the membrane mass now will be able to travel further outwards over a given amount of time. It is worth noting that fluctuations in the direction of outgrowth activity could occur as very rapid minute movements of membrane mass. When observed at the macro-scale, these micro-scale fluctuations might not be seen. Instead, the outward movement of an extension will appear as linear, straight-line, movement. In this paper, “fluctuation” refers to variation in the direction of outgrowth activity. “Outgrowth” refers to the movement of membrane mass at the micro-scale, whereas “extension” refers to the movement of the axon that is observed at the macro-scale.

The SDAL model makes distinctive predictions about UNC-40-mediated outgrowth activity *in vivo* and the direction of extension. In a deterministic model, the direction of extension is determined by the outward movement of the membrane from the site where UNC-6 and UNC 40 interact. Positional information is encoded by gradients so that UNC-6 guides extension towards the UNC-6 source. In the SDAL model, UNC-6 and other extracellular cues govern the probability of UNC-40 asymmetric localization, and subsequent UNC-40-mediated outgrowth, at each surface of the membrane (XU *et al.* 2009; KULKARNI *et al.* 2013). The direction of outgrowth is determined by a directional bias that is created over time by the combined effect of extracellular cues. If, for example, the probability of outgrowth towards a ventral UNC-6 source is 0.3 and the probability of dorsal outgrowth is 0.3 and of anterior outgrowth is 0.4, the direction of outgrowth will be in the anterior direction. That is, at any instance of time there is a chance that outgrowth movement will be directed ventrally towards the UNC-6 source, however over a longer period of time the outgrowth will travel anteriorly because there is always a greater likelihood that outgrowth will be anterior instead of ventral or dorsal. To reiterate, the direction of extension is a product of a stochastic process, in which the outcome evolves over time. The probability of ventral outgrowth created by the UNC-40-mediated response to UNC-6 is required for the anterior bias. Without the ventrally directed outgrowth in response to UNC-6, the probability of ventral outgrowth would decrease, shifting the directional bias. Thus, when considered as a stochastic process, the observed directional response to the interactions between UNC-40 and UNC-6 is not necessarily extension towards the UNC-6 source. Because of SDAL, positional information is encoded by the location and level of the extracellular cues along the surface of the neuron.

The SDAL model suggested that UNC-40 activity could affect extension movement in ways that had not been obvious. The first insight is that forward movement of an extension could be inhibited as it moves towards a source of a cue that promotes outgrowth (Figure 3A). At the leading edge of an outgrowth, a strong directional basis for movement towards an UNC-6 source occurs only as long as the probability of UNC-40 localization at surfaces facing towards the source are greater than the probability of UNC-40 localization at surfaces facing other directions. As an extension moves towards an UNC-6 source a higher proportion of the UNC-40 receptors that flank the leading edge can become ligated (Figure 3B). Because of the SDAL process, this will increase the probability of localization and outgrowth at the flanking sites while decreasing the probability of localization and outgrowth at the leading edge. The result will be an increase in random movement and a decrease in the outward displacement of the membrane’s mass. Paradoxically, the rate of extension will decrease as the extension moves towards the UNC-6 source (Figure 3C). It is worth noting that even if the probability of outgrowth in each direction becomes equal, there will still be a directional bias. For example, if the probability of outgrowth towards a ventral UNC-6 source is 0.33 and of anterior outgrowth is 0.33 and of posterior outgrowth is 0.33, the directional bias is ventral. (The probability of movement in the direction of the axon shaft (backwards) is low.)

**Figure 3.**
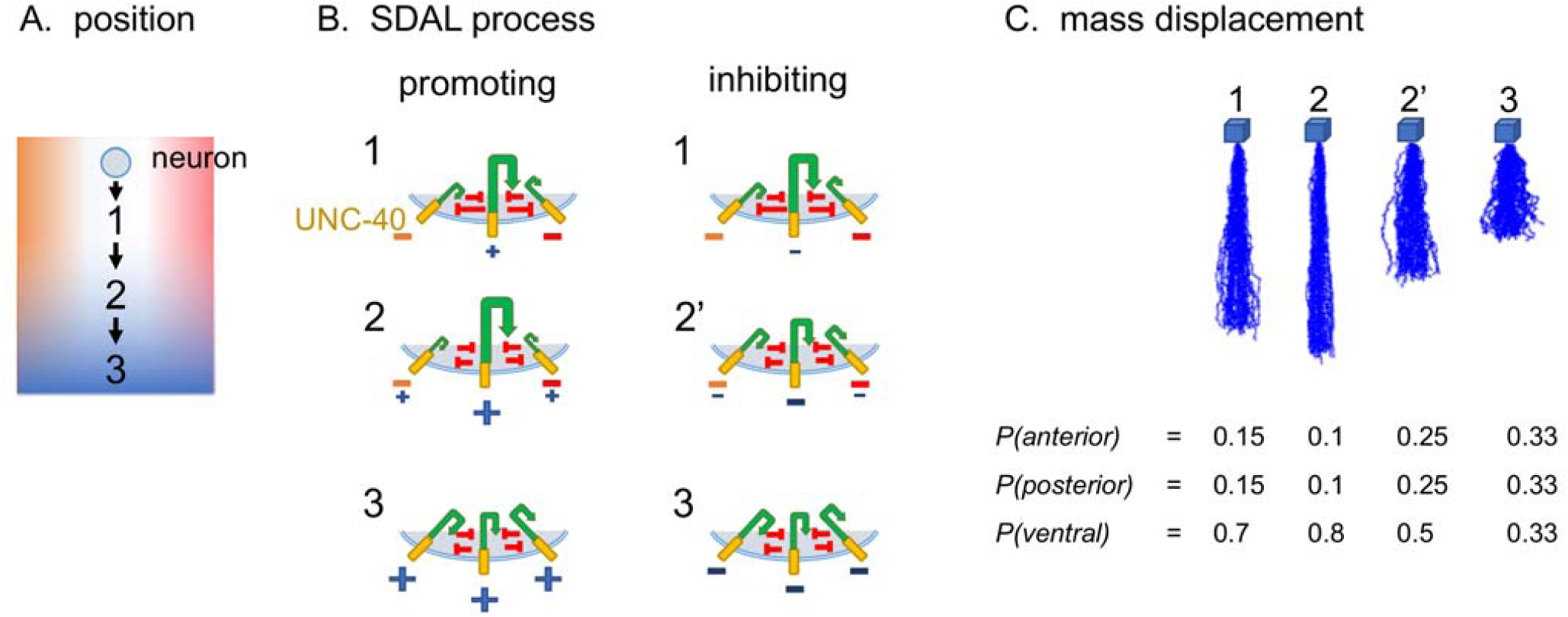
Model for outgrowth movement towards extracellular cues that promote or inhibit outgrowth activity. **(A)** Schematic diagram of the outgrowth of a neuron through an environment of multiple extracellular cues. These cues may be molecules present at the surfaces of surrounding cells and extracellular matrix, or they may be physical interactions that influence outgrowth activity. The extracellular cues are represented as color gradients of blue, orange, and red. The neuron’s response to cues arranged along the anterior/posterior axis (orange and red), create an equal probability for UNC-40 asymmetric localization and outgrowth in the anterior and posterior directions. The extension transverses three different positions (1-3) as it develops towards a ventral source of a cue (blue). **(B)** The SDAL process is illustrated as in Figure 1 for the three positions shown in A. Shown are scenarios for movement towards a cue (A, blue) that promotes outgrowth (blue +) or that inhibits outgrowth (blue -). At positions 1 and 2 cues along the anterior/posterior axis (orange - and red -) prevent outgrowth in the anterior or posterior directions. At position 3, the cue from the ventral source predominates. **(C)** Random walk modeling as described in Figure 2E. At each position (A, 1-3), cues alter the probability distribution for the direction of localization and outgrowth. Below each plot is the probability distribution used to create the random walk (see Materials and Methods). Probability distributions were selected to represent how different levels of the ventral cue might change the probability distribution at each position. The plots illustrate the probability density function of the position of mass as a function of space and time if movement occurred according to that probability distribution. For both scenarios, an equal probability of anterior and posterior outgrowth can allow a ventral directional bias at position 1. Movement towards a promoting cue source can allow a greater probability for ventral outgrowth and, correspondingly, a lower probability for anterior and posterior outgrowth (position 2). Movement towards an inhibiting cue source can allow a lower probability for ventral outgrowth and, correspondingly, a greater probability for anterior and posterior outgrowth (position 2’). The ventral direction bias is maintained. In either scenario, an equal probability for outgrowth in all directions may occur as the receptors become saturated because of the high level of the cue from the ventral source (position 3). The modeling predicts that in both scenarios changing levels of the ventral cue will not alter the direction of outward movement, although it may alter the outward displacement of the membrane’s mass. See text for details.

A second insight is that an extension could move towards the source of a cue that inhibits outgrowth (Figure 3A). For example, if together the extracellular cues create a probability for ventral outgrowth of 0.7, for anterior outgrowth of 0.15, and for posterior outgrowth of 0.15, a directional bias for ventral outgrowth is created (Figure 3B). This can occur even if there is a ventral source of an inhibitory cue. The extension can move ventrally towards this inhibitory cue source. Eventually the probability for ventral outgrowth might change to 0.33, anterior to 0.33, and posterior to 0.33 (Figure 3C). But even in this case, there is still a directional bias for ventral outgrowth and extension will continue to move towards the source of the inhibitory cue.

The model predicts that movement towards a source of a cue causes the system to trend towards a state where the probabilities of outgrowth in different directions become equal. Axons often change their trajectory near the source of a cue. It is possible that the state is important because the equilibrium might allow cues to be more effectual at reorienting outgrowth (Figure 4).

**Figure 4.**
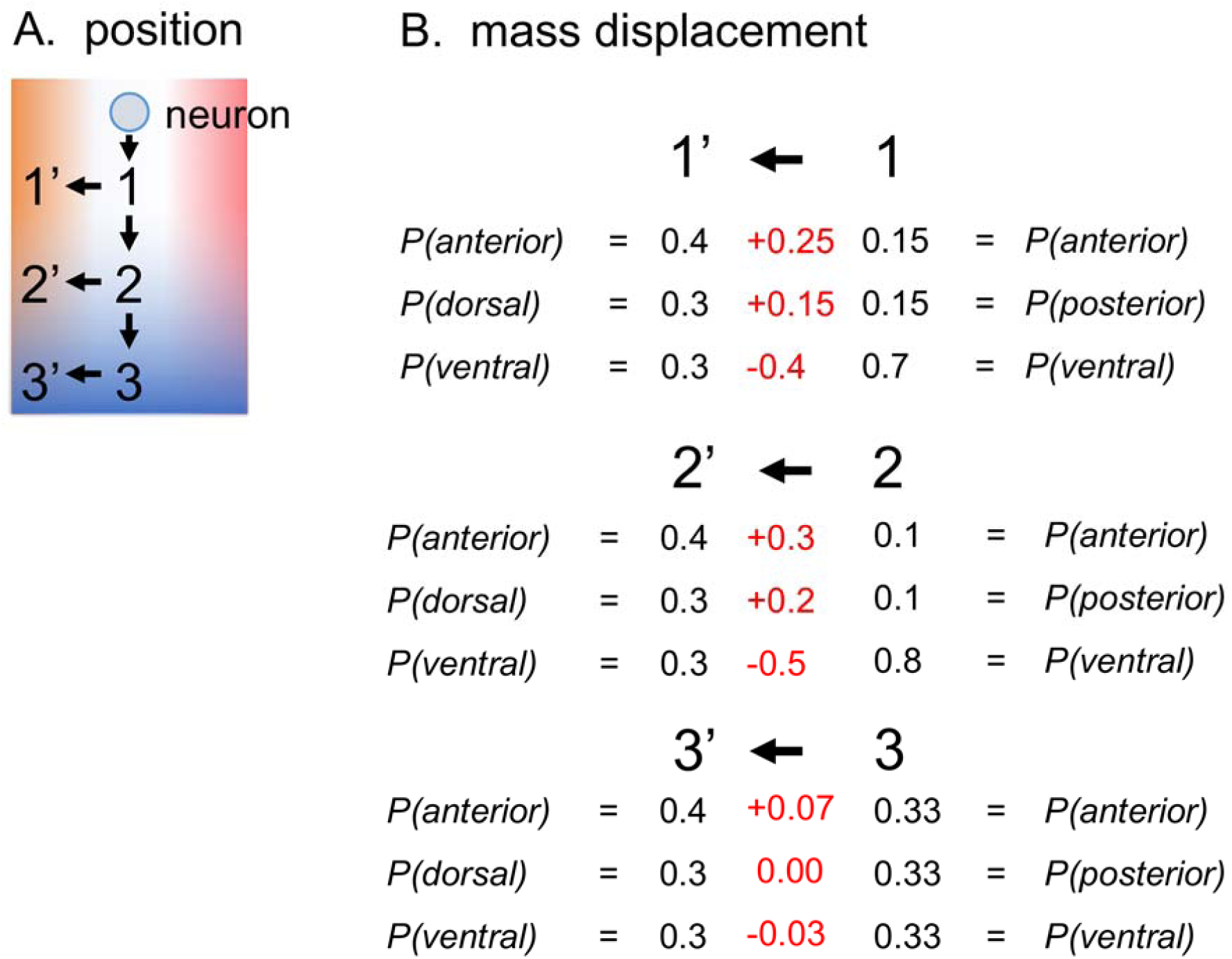
Model for outgrowth movement that changes direction. **(A)** Schematic diagram of the outgrowth of a neuron through an environment of multiple extracellular cues as described in Figure 3A. Positions 1’-3’ represent the position after a change from ventral to anterior outgrowth. **(B)** At each position (A, 1-3), the probability distribution for the direction of localization and outgrowth is given as in Figure 3C. In order for the direction of outgrowth to shift anteriorly at each position, the probability distribution must shift to create a bias for anteriorly directed outgrowth. This is depicted by the probability distribution of 0.4 anterior, 0.3 dorsal, and 0.3 ventral for positions 1’-3’. Numbers in red indicate the degree to which the probabilities must change between the positions. The model predicts that as the system trends towards a state where the probabilities of outgrowth in different directions become equal, cues that could shift the direction bias become more effectual.

The third insight is that multiple extensions from a neuron could move in the same direction without having to follow prepatterned extracellular pathways. Some neurons send out multiple extensions that run in parallel towards a target. It is commonly proposed that these patterns form because extensions follow parallel pathways that were previously formed by extracellular guidance cues. The SDAL model suggests that multiple UNC-40-mediated outgrowths can be initiated at a leading surface and that multiple extensions can maintain their positions without having to follow prepatterned extracellular pathways. In this model, a separate extension begins to form at the leading edge because the directional bias at one site becomes greater than that at flanking sites. We propose that along the leading edge the self-organizing UNC-40 localization process can create multiple sites that have a greater directional bias (Figure 5A). The positive and negative feedback loops of the SDAL process can allow spatial patterns of outgrowth to develop autonomously. Once these sites are established, outgrowth can proceed from each site in the same direction (Figure 5B). The strongest directional bias is created when the probabilities for outgrowth are equal in the directions perpendicular to the direction of extension. The actual value of the perpendicular probabilities is not crucial for establishing a directional bias. Even though the value of the perpendicular probabilities may vary depending on the position of outgrowth along the perpendicular axis, the direction of outgrowth will be the same. If a perpendicular equilibrium is maintained, then cues that affect UNC-40 localization and outgrowth and which are distributed along the perpendicular axis will have little effect on the direction of outgrowth. Such an equilibrium can be established if the outgrowth effects of cues distributed along the perpendicular axis balance out each other. Such a condition could be established by cues that effect outgrowth equally at surfaces facing the perpendicular axis. Even if cues are distributed in a gradient an equilibrium could exist. Studies indicate that gradient steepness, rather than the concentration of cues, is important for growth cone turning and guidance (BAIER AND BONHOEFFER 1992; ROSOFF *et al.* 2004; MORTIMER *et al.* 2010; SLOAN *et al.* 2015). Therefore, cues may create a perpendicular equilibrium if they are distributed in a shallow gradient along the perpendicular axis.

**Figure 5.**
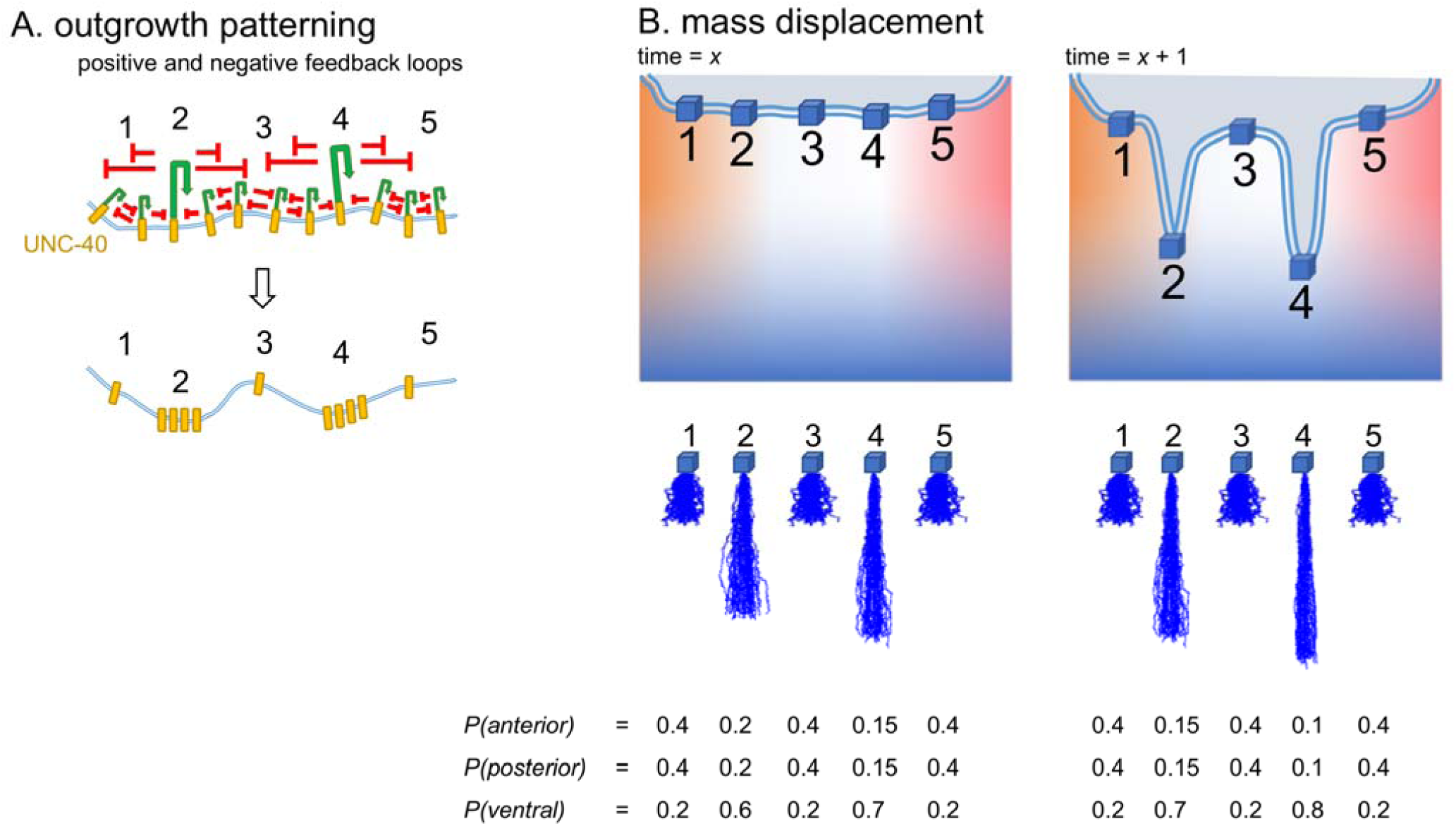
Model for the development of multiple outgrowths that extend in the same direction. **(A)** The SDAL process is illustrated for sites along a surface of a neuron as in Figure 1. The positive and negative feedback loops of the SDAL process allow spatial patterns of outgrowth to develop autonomously. The number of sites where a strong directional bias is ultimately created is dictated by the relative effectiveness of the positive and negative feedback loops. **(B)** Schematic diagram of the outgrowth of a neuron through an environment of multiple extracellular cues as described in Figure 3A. The flow of membrane mass (box) at different sites depends on the probability distribution for the direction of outgrowth created at regions along the surface. Random walk modeling as described in Figure 2E is shown below the schematic diagram. At time X, two sites which have a greater directional bias (2 and 4) are established by the SDAL process as depicted in A. Cues may not be present in steep gradients along the axis perpendicular to the direction of extension. The response to these cues creates probabilities for outgrowth that are equal in the perpendicular directions. The greatest directional bias is created when there is an equilibrium for the probability of outgrowth in perpendicular directions. Because cue levels may vary gradually along the perpendicular axis, the strength of the directional bias at sites may differ, however the bias will be oriented in the same direction. At time *x* + 1, positions 2 and 4 have proceeded further outward because of greater membrane displacement. This effect is magnified by increasing levels of the outgrowth promoting cue (blue). This activity together, when averaged over time across a surface, is predicted to cause the dynamic development of multiple extensions.

A fourth insight is that cues can direct movement without being in a concentration gradient. The SDAL activity within the cell initiates random walk movement. As long as an equilibrium along the perpendicular axis exists, a directional bias along the other axis can be created. In Figure 5B outgrowth is towards a ventral cue source, and as outgrowth moves up the concentration gradient of this cue the probability of outgrowth in each direction changes. However, movement towards the source would still occur if the concentration of extracellular cues remains constant and the probabilities never change. Because of the SDAL process, a directional bias can be maintained along a track of a uniformly distributed cue.

The last insight is that extracellular cues could affect UNC-40 localization and outgrowth, but not affect the direction of outgrowth. However, these cues could have an effect on the morphology and patterning of an extension. A candidate for such a cue is EGL-20 (wnt). The *egl-20* gene is one of several Wnt genes in *C. elegans.* These genes are expressed in a series of partially overlapping domains along the anteroposterior axis of the animal (SAWA AND KORSWAGEN 2013). EGL-20 is expressed in cells posterior to HSN (WHANGBO AND KENYON 1999; PAN *et al.* 2006; HARTERINK *et al.* 2011). The sources of UNC-6 and EGL-20 are roughly perpendicular to each other. We have observed that loss of EGL-20 function causes UNC-40 asymmetrical localization to orient to randomly selected surfaces of HSN and causes the axon to initially extend from the HSN cell body in different directions (KULKARNI *et al.* 2013; TANG AND WADSWORTH 2014). UNC-6 and EGL-20 signaling could both impinge on the feedback loops that regulate UNC-40 SDAL. In doing so, these cues would act together to influence the pattern of extension. In this paper, we provide further genetic evidence that the downstream signals from both cues converge to regulate the UNC-40 SDAL process.

We suggest that UNC-5 plays an important role in coordinating the UNC-40 SDAL process with non-UNC-40-mediated responses that affect outgrowth. Previously we reported that loss of UNC-5 causes UNC-40 asymmetrical localization to orient to randomly selected surfaces of HSN, causing the axon to initially extend in different directions (KULKARNI *et al.* 2013). This suggests that UNC-5 functions to increase the probability of UNC-40 asymmetric localization being oriented to the site of UNC-6 and UNC-40 interaction. That is, UNC-5 promotes straight-line motion by inhibiting the degree to which the direction of UNC-40-mediated outgrowth fluctuates. UNC-5 has other functions as well. UNC-5 is primarily known for its role in mediating movement away from UNC-6 sources. For example, UNC-5 is required for the dorsal migration of DA and DB motor neuron axons away from ventral UNC-6 sources (HEDGECOCK *et al.* 1990). DA and DB guidance utilizes both UNC-40-dependent and UNC-40-independent pathways, although guidance is significantly less disrupted by loss of UNC-40 than by loss of UNC-5 (HEDGECOCK *et al.* 1990; MACNEIL *et al.* 2009). In keeping with the SDAL model, we predict that UNC-5 increases the probability of non-UNC-40-mediated outgrowth being oriented towards sites where there are not interactions with UNC-6. This increases the probability of outgrowth movement in directions not towards UNC-6 sources. Finally, we predict that UNC-5 function can also increase the probability that non-UNC-40-mediated outgrowth will orient to the site of interaction between non-UNC-40 receptors and non-UNC-6 extracellular cues. Evidence for this comes from the observation that in *rpm-1* mutants the overextension of the PLM axon can be suppressed by loss of UNC-5, but not by the loss of UNC 40 or UNC-6 (LI *et al.* 2008). In summary, we believe UNC-5 can: 1) increase the probability of UNC-40-mediated outgrowth at the sites of UNC-6 and UNC-40 interactions; 2) decrease the probability of UNC-40-mediated outgrowth at sites where UNC-6 is not present; 3) decrease the probability of non-UNC-40-mediated outgrowth at the site of UNC-6 and UNC-40 interactions; and 4) increase the probability of non-UNC-40-mediated outgrowth at sites where there are no UNC-6 interactions. These functions can be considered in terms of the positive and negative feedback loops of the SDAL model (Figure 6), where UNC-5 helps regulated the feedback loops associated with UNC-40 activity.

**Figure 6.**
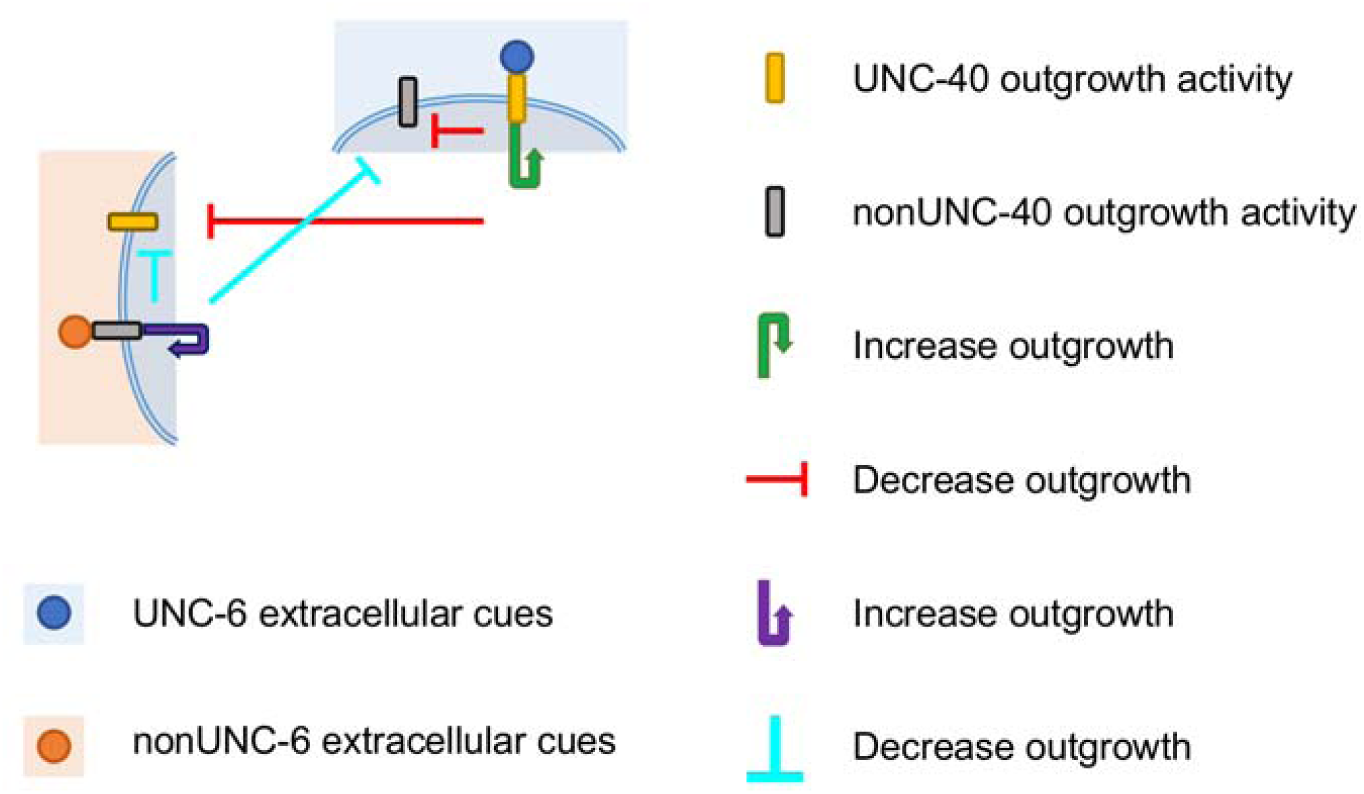
Model for the control of outgrowth activity by SDAL. Schematic diagram of the control of UNC-40- and nonUNC-40-mediated outgrowth activity. Neuronal surfaces of the neuron are subjected to different levels of UNC-6 (blue) as well as nonUNC-6 extracellular cues (orange). The SDAL process regulates both UNC-40 and nonUNC-40-mediated outgrowth activity. Positive feedback (arrows) amplifies the polarized response to an extracellular cue, while negative feedback (lines) limits the response and can confine the positive feedback to the site of ligand interaction. The long-range negative feedback mediated by UNC-40 inhibits the UNC-40 response to UNC-6, as well as nonUNC-40 activity. Similarly, long-range negative feedback mediated by nonUNC-40 activity inhibits the UNC-40 response to UNC-6.

Because of these ideas, we reasoned that UNC-5 activity could affect extension movement in ways that had not been obvious to us. First, UNC-5 might affect the patterning of extension that travels *towards* an UNC-6 source. As discussed above, previous evidence suggests UNC-5 regulates the asymmetric localization of UNC-40. UNC-5 interactions with UNC-6 and UNC-40 could influence the feedback loops (Figures 1 and 6). By regulating the degree to which the direction of UNC-40-mediated outgrowth fluctuates, UNC-5 could affect random movement and the outward displacement of membrane mass. This could affect the rate of extension towards an UNC-6 source or whether extension can occur. In cases where multiple extensions form from a surface, the effect UNC-5 has on the loops could influence whether sites with a predominate directional bias can be established.

Second, UNC-5 could play a role in determining whether an extension changes direction. As discussed earlier, if the UNC-40 receptors become saturated near an UNC-6 source then the probability of UNC-40-mediated outgrowth towards the source and along the perpendicular axis tends to become equal. At this point, even a small increase in the probability of non-UNC 40-mediated outgrowth to the site of non-UNC-6 and non-UNC-40 interactions could alter the directional bias (Figure 4). A change in UNC-5 activity could help promote a shift from a directional bias determined by UNC-40 and UNC-6 interactions, to one determined by non UNC-40 and non-UNC-6 interaction.

Because of the predictions that the UNC-40 SDAL model makes, we decided to reexamine the *unc-5* loss-of-function phenotypes and to investigate genetic interactions among *unc-5, unc-6, unc-40* and *egl-20* that regulated the asymmetric localization of UNC-40. We find evidence that UNC-5 regulates the length and number of processes that extend towards an UNC-6 source and that UNC-5 helps control the ability of axons to extend in different directions. In addition, we find genetic interactions that suggest UNC-5, together with UNC-53 (NAV2), functions to regulate UNC-40 SDAL in response to the UNC-6 and EGL-20 (wnt) extracellular cues. We suggest that the SDAL model is useful for understanding how genes regulate the patterning of axon extensions.

## Materials and Methods

### Strains

Strains were handled at 20 °C using standard methods (Brenner, 1974) unless stated otherwise. A Bristol strain N2 was used as wild type. The following alleles were used: **LGI**, *unc-40(e1430), unc 40(ur304), zdIs5[mec-4::GFP]*; **LGII**, *unc-53(n152)*; **LGIV**, *unc-5(e152), unc-5(e53), unc-5(ev480), unc 5(ev585),egl-20(n585), kyIs262[unc-86::myr-GFP;odr-1::dsRed];* **LGIV**, *madd-2(ky592), madd-2(tr103)*; **LGX**, *mig-15(rh148), unc-6(ev400)*, *sax-3(ky123), sax-3(ky200).* Transgenes maintained as extrachromosomal arrays included: *kyEx1212 [unc-86::unc-40-GFP;odr 1::dsRed].*

### Analysis of axon outgrowth and cell body position

HSN neurons were visualized using expression of the transgene *kyIs262[unc-86::myr-GFP].* The mechanosensory neurons, AVM, ALM, and PLM, were visualized using the expression of the transgene *zdIs5[Pmec-4::GFP]*. Synchronized worms were obtained by allowing eggs to hatch overnight in M9 buffer without food. The larval stage was determined by using differential interference contrast (DIC) microscopy to examine the gonad cell number and the gonad size. Staged larvae were mounted on a 5% agarose pad with 10 mM levamisole buffer. Images were taken using epifluorescent microscopy with a Zeiss 63X water immersion objective.

The number of processes during early L1 larval stage was scored by counting the number of processes that extended for a distance greater than the length of one cell body. We report instances in which there were no such processes, one process or more than one processes. In the L2 larval stage, a single early process was scored if there was only one major extension from the ventral leading edge. The HSN cell body in L2 stage larvae was scored as dorsal if the cell body had failed to migrate ventrally and was not positioned near the PLM axon. In L4 stage larvae, a multiple ventral processes phenotype was scored if more than one major extension protruded from the ventral side of cell body.

Extension into the nerve ring was scored as defective if the axon did not extend further than approximately half the width of the nerve ring. Anterior extension was scored as defective if the axon did not extend further anteriorly than the nerve ring. PLM axons are scored as over-extending if they extended further anterior than the position of the ALM cell body.

### Analysis of the direction of HSN outgrowth

HSN was visualized using the transgene *kyIs262[unc-86::myr-GFP].* L4 stage larvae were mounted on a 5% agarose pad with 10 mM levamisole buffer. An anterior protrusion was scored if the axon extended from the anterior side of the cell body for a distance greater than the length of three cell bodies. A dorsal or posterior protrusion was scored if the axon extended dorsally or posteriorly for a distance greater than two cell body lengths. HSN was considered multipolar if more than one process extended a length greater than one cell body. Images were taken using epifluorescent microscopy with a Zeiss 40X objective.

### Analysis of the UNC-40::GFP localization in L2 stage animal

For analysis of UNC-40::GFP localization, L2 stage larvae with the transgenic marker *kyEx1212[unc-86::unc-40::GFP; odr-1::dsRed]* were mounted on a 5% agarose pad with 10 mM levamisole buffer. Staging was determined by examining the gonad cell number and the gonad size under differential interference contrast (DIC) microscopy. Images were taken using epifluorescent microscopy with a Zeiss 63X water immersion objective. The UNC-40::GFP localization was determined by measuring the average intensity under lines drawn along the dorsal and ventral edges of each HSN cell body by using ImageJ software. For analysis of the anterior–posterior orientation of UNC-40::GFP, the dorsal segment was geometrically divided into three equal lengths (dorsal anterior, dorsal central and dorsal posterior segments). The line-scan intensity plots of each of these segments were recorded. ANOVA test was used to determine if there is a significant difference between intensities of the three segments. The dorsal distribution was considered uniform if p≥0.05 and was considered asymmetrical if p≤0.05. Within an asymmetric population, the highest percent intensity was considered to localize UNC-40::GFP to either anterior, posterior or central domain of the dorsal surface.

### Computations

A program to simulate a two-dimensional lattice random walk based on the probability of dorsal, ventral, anterior, and posterior outgrowth for a mutant (Table 1) was created using MATLAB. (The directions of the axons from multipolar neurons were not scored. These axons appear to behave in the same manner as the axons from monopolar neurons, but this has not yet been tested.) The probability of dorsal, ventral, anterior, or posterior outgrowth was assigned for the direction of each step of a random walk moving up, down, left or right, respectively (Figure 9). Each variable is considered independent and identically distributed. Simulations of 500 equal size steps (size =1) were plotted for 50 tracks (Figure 1B, 5B and 6B inserts). A Gaussian distribution for the final positions of the tracks was generated using Matlab’s random function (Figure 6).

**Table 1.** Direction of Axon Formation from the HSN Cell Body

| Table 1. Direction of Axon Formation from the HSN Cell Body |  |  |  |  |  |  |  |
| --- | --- | --- | --- | --- | --- | --- | --- |
|  | direction of axon protrusion |  |  |  |  | *n* | reference |
|  | dorsal | ventral | anterior | posterior | multipolar |  |  |
|  | % | % | % | % | % |  |  |
| wildtype | 0 | 96±2 | 3±2 | 0 | 1±1 | 221 | (KULKARNI *et al.* 2013) |
| *unc-6(ev400)* | 2±2 | 3±2 | 81±2 | 8±2 | 6±1 | 218 | (KULKARNI *et al.* 2013) |
| *unc-40(e1430)* | 2±1 | 6±2 | 67±2 | 19±1 | 6±1 | 183 | (KULKARNI *et al.* 2013) |
| *unc-5(e53)* | 0 | 75±3 | 19±2 | 1±1 | 5±1 | 245 | (YANG *et al.* 2014) |
| *unc-53(n152)* | 0 | 67±3 | 22±2 | 5±1 | 6±1 | 238 | (KULKARNI *et al.* 2013) |
| *sax-3(ky123)* | 2±1 | 31±1 | 21±1 | 37±2 | 9±2 | 232 | (TANG AND WADSWORTH 2014) |
| *sax-3(ky200)\** | 2±1 | 32±1 | 19±2 | 42±3 | 5±2 | 198 | (TANG AND WADSWORTH 2014) |
| *unc-5(e53);sax-3(ky200)* | 2±1 | 40±3 | 24±2 | 28±2 | 6±1 | 120 |  |
| *unc-5(e53);unc-6(ev400)* | 4±2 | 5±3 | 59±4 | 22±4 | 9±1 | 201 |  |
| *unc-5(e53);egl-20(n585)* | 3±1 | 28±4 | 22±4 | 35±5 | 11±2 | 114 |  |
| *unc-53(n152);unc-5(e53)* | 0 | 19±1 | 62±2 | 17±1 | 3±1 | 224 | (KULKARNI *et al.* 2013) |
| *unc-53(n152);unc-6(ev400)* | 24±2 | 0 | 19±2 | 22±2 | 34±3 | 144 | (KULKARNI *et al.* 2013) |
| *unc-53(n152);sax-3(ky123)* | 1±1 | 47±3 | 24±2 | 23±5 | 6±3 | 207 | (TANG AND WADSWORTH 2014) |
| *unc-40(e1430);unc-5(e53)* | 5±1 | 6±1 | 55±2 | 19±2 | 14±1 | 196 | (KULKARNI *et al.* 2013) |
| *unc-40(e1430);sax-3(ky200)\** | 14±3 | 2±1 | 40±2 | 35±3 | 9±4 | 191 | (TANG AND WADSWORTH 2014) |
| *sax-3(ky200)\*; unc-6(ev400)* | 8±1 | 8±2 | 49±3 | 20±5 | 14±2 | 211 | (TANG AND WADSWORTH 2014) |
| *unc-53(n152);unc-5(e53);unc-6(ev400)* | 23±2 | 0 | 34±2 | 15±2 | 28±2 | 148 | (KULKARNI *et al.* 2013) |
| *unc-53(n152);sax-3(ky200)\*;unc-6(ev400)* | 11±2 | 2±1 | 33±4 | 30±3 | 25±5 | 189 |  |
| *egl-20(n585)* | 0 | 64±2 | 21±2 | 7±1 | 8±1 | 304 | (TANG AND WADSWORTH 2014) |
| *egl-20(n585); unc-6(ev400)* | 18±2 | 0 | 43±2 | 15±2 | 24±2 | 205 | (TANG AND WADSWORTH 2014) |
| *unc-40(e1430); egl-20(n585)* | 6±2 | 17±2 | 45±5 | 15±2 | 16±2 | 173 | (TANG AND WADSWORTH 2014) |
| *egl-20(n585);sax-3(ky123)* | 1±1 | 12±2 | 39±2 | 39±1 | 8±3 | 177 | (TANG AND WADSWORTH 2014) |
| *madd-2(tr103)* | 0 | 19±2 | 55±5 | 17±4 | 8±2 | 179 |  |
| *madd-2(ky592)* | 0 | 52±2 | 43±2 | 5±1 | 0 | 95 |  |
| *unc-5(e53);madd-2(tr103)* | 3±1 | 15±2 | 52±4 | 17±4 | 13±1 | 197 |  |
| *madd-2(tr103);sax-3(ky123)* | 2 | 24±3 | 19±4 | 47±1 | 7±2 | 171 |  |
| *unc-53(n152);madd-2(tr103)* | 1±1 | 15±2 | 43±2 | 17±1 | 24±4 | 148 |  |
| *mig-15(rh326)* | 2±1 | 15±1 | 24±3 | 11±3 | 48±8 | 131 | (YANG *et al.* 2014) |

Numbers represent percentage value ± SEM.
\*Animals grown at the *sax-3(ky200)* restrictive temperature (25°C).

The mean squared displacement (MSD) is used to provide a quantitative characteristic of the motion that would be created by the outgrowth activity undergoing the random walk. Using the random walks generated for a mutant the MSD can be calculated:

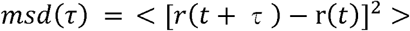

Here, r(*t*) is the position at time *t* and τ is the lag time between two positions used to calculate the displacement, Δr(τ) = r(*t*+τ) - r(*t*). The time-average over *t* and the ensemble-average over the 50 trajectories were calculated. This yields the MSD as a function of the lag time. A coefficient giving the relative rate of diffusion was derived from a linear fit of the curve. The first two lag time points were not considered, as the paths often approximate a straight line at short intervals.

## Results

### UNC-5 regulates the pattern of outgrowth from the HSN neuron

To investigate whether UNC-5 activity can regulate the length or number of processes that a neuron can develop when outgrowth is towards an UNC-6 source, we examined the development of the HSN axon in *unc-5* mutations. The HSN neuron sends a single axon to the ventral nerve cord, which is a source of the UNC-6 cue (WADSWORTH *et al.* 1996; ADLER *et al.* 2006; ASAKURA *et al.* 2007). Axon formation is dynamic (ADLER *et al.* 2006). Shortly after hatching, HSN extends short neurites in different directions. These neurites, which dynamically extend and retract filopodia, become restricted to the ventral side of the neuron where a leading edge forms. Multiple neurites extend from this surface until one develops into a single axon extending to the ventral nerve cord. Measurements of growth cone size, maximal length, and duration of growth cone filopodia indicate that UNC-6, UNC-40, and UNC-5 control the dynamics of protrusion (NORRIS AND LUNDQUIST 2011).

We observe that in *unc-5* mutants, the patterns of extension are altered. In wild-type animals at the L1 stage of development most HSN neurons extends more than one short neurite, however in *unc 5(e53)* mutants nearly half the neurons do not extend a process (Figures 7A and 7B). During the L2 stage in wild-type animals a prominent ventral leading edge forms and the cell body undergoes a short ventral migration that is completed by the L3 stage. By comparison, in *unc-5* mutants the cell body may fail to migrate and instead a single large ventral process may form early during the L2 stage (Figures 7A, 7C and 7E). It may be that the ventral migration of the HSN cell body requires the development of a large leading edge with multiple extensions. Together the observations indicate that loss of *unc-5* function affects the patterning of outgrowth, *i.e.* the timing, length, and number of extensions that form. Loss of *unc-5* function does not prevent movement, in fact, a single large ventral extension can form in the mutant at a time that is even earlier than when a single ventral extension can be observed in wildtype. The earlier appearance of a single ventral extension in *unc-5* mutants appears to be the result of a difference in morphology, rather than of developmental timing. The failure of the HSN cell body to migrate ventrally and the different pattern of outgrowth at the leading edge causes an earlier discernable single extension.

**Figure 7.**
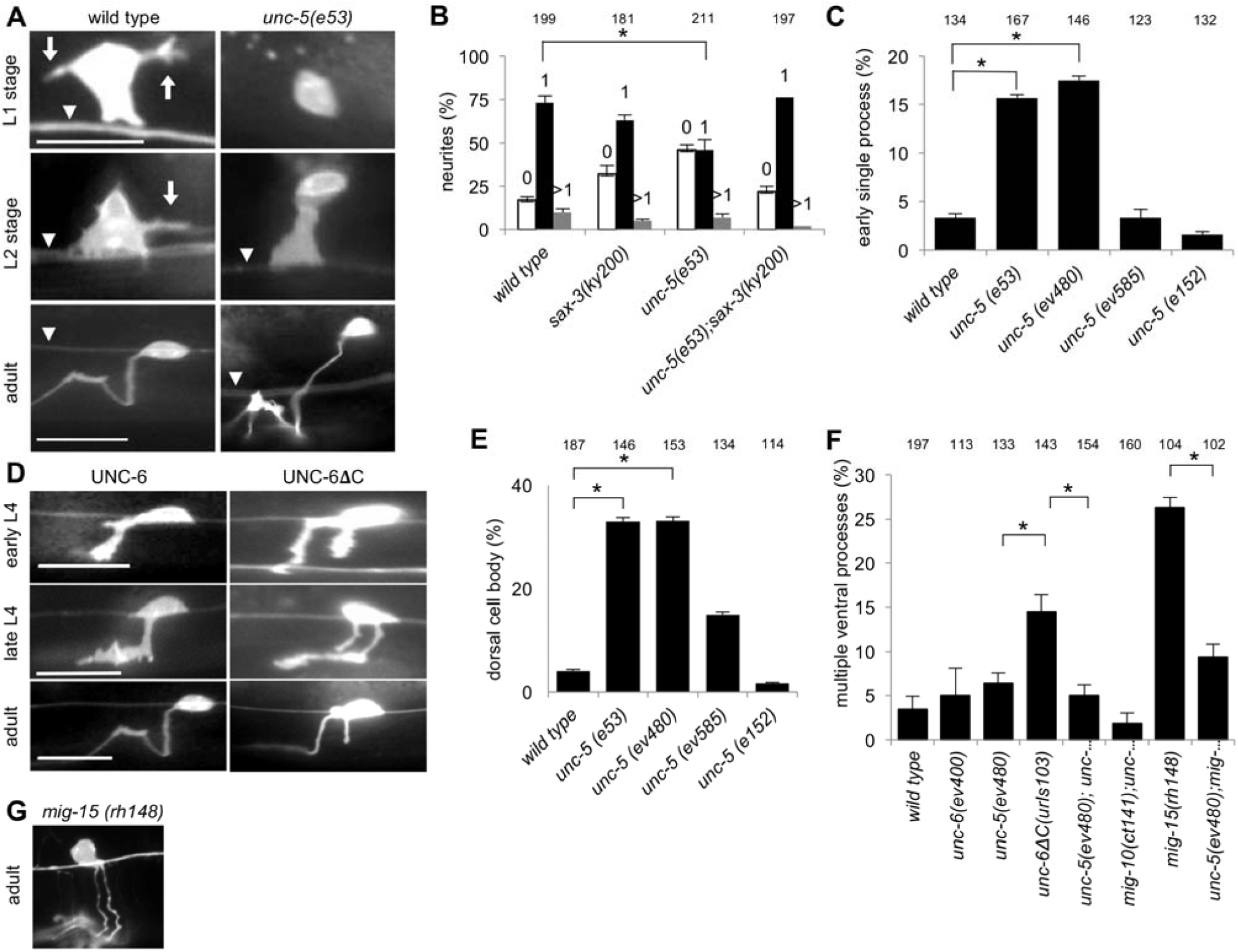
UNC-5 regulates the patterning of outgrowth extensions from HSN. **(A)** Photomicrographs of HSN at the L1, L2, and adult stages in wildtype and *unc-5(e53)* mutants. In L1 and L2 animals neurite extensions (arrows) are often observed in wild-type animals but are more rare in *unc-5* mutants. The short ventral migration of the cell body that occurs in wild-type animal sometimes fails in *unc-5* mutants, leaving the cell body farther from the PLM axon (arrowhead) with a single longer ventral extension. The position of the cell body remains dorsal. Scale bar: 10 μm. **(B)** The percentage of HSN neuron with 0, 1, or more than 1 neurite extension at the L1 stage. In *unc-5* mutants nearly half of the neurons do not extend a process. Error bars indicated the standard error mean; n values are indicated above each column. Significant differences (two-tailed t-test), *P<0.001. **(C)** The percentage of HSN neurons with a single long extension at the L2 stage. Several *unc-5* alleles were tested as described in the text. In mutants with loss-of-function there is more often a single extension from the cell body and the cell body is dorsally mispositioned. **(D)** Photomicrographs of HSN at the early L4, late L4, and adult stages in wildtype and in animals expressing UNC-6ΔC. The expression of UNC-6ΔC induces multiple processes, most often two major extensions, that are guided ventrally. **(E)** The percentage of HSN neurons with a cell body mispositioned dorsally at the L2 stage. In loss-of-function mutants the cell body often fails to undertake a short ventral migration during the L2 stage. The migration is not delayed, but rather it remains dorsal. **(F)** The percentage of HSN neurons with multiple ventral extensions at the L4 stage. The additional processes induced by UNC-6ΔC can be suppressed by *unc-5* and *mig-10* mutations. Additional processes induced by *mig-15(rh148)* can also be suppressed by the *unc-5* mutation. **(G)** Photomicrographs of HSN at adult stages in a *mig-15* mutant. Similar to UNC-6ΔC expression, *mig-15* mutations can also cause additional processes that are guided ventrally (YANG *et al.* 2014).

We tested four different *unc-5* alleles in these experiments. The *unc-5(e53)* allele is a putative molecular null allele, *unc-5(ev480)* is predicted to truncate UNC-5 after the cytoplasmic ZU-5 domain and before the Death Domain, *unc-5(e152)* is predicted to truncate UNC-5 before the ZU-5 domain and Death Domain, and *unc-5(ev585)* is a missense allele that affects a predicted disulfide bond in the extracellular Ig(C) domain (KILLEEN *et al.* 2002). Although both the *unc 5(ev480)* and *unc-5(e152)* are predicted to cause premature termination of protein translation in the cytodomain, the *unc-5(e152)* product retains the signaling activity that prevents these phenotypes. Based on other phenotypes, previous studies reported that the *unc-5(e152)* allele retains UNC-40-dependent signaling functions (MERZ *et al.* 2001; KILLEEN *et al.* 2002).

### UNC-5 is required for the induction of multiple HSN axons by UNC-6ΔC and a mig-15 mutation

The results above suggest that UNC-5 activity can regulate the number of HSN extensions that form. To further test this hypothesis, we checked whether loss of UNC-5 function can suppress the development of additional processes that can be induced. Previously we reported that expression of the N-terminal fragment of UNC-6, UNC-6ΔC, induces excessive branching of ventral nerve cord motor neurons and that loss of UNC-5 function can suppress this branching (LIM *et al.* 1999). We now report that HSN develops an extra process in response to UNC-6ΔC and that loss of UNC-5 function suppresses the development of this extra process (Figures 7D and 7F).

To investigate whether this UNC-5 activity might involve known effectors of asymmetric neuronal outgrowth, we tested for genetic interactions between *unc-5* and both *mig-10* and *mig-15.* MIG-10 (lamellipodin) is a cytoplasmic adaptor protein that can act cell-autonomously to promote UNC-40 mediated asymmetric outgrowth (ADLER *et al.* 2006; CHANG *et al.* 2006; QUINN *et al.* 2006; QUINN *et al.* 2008; MCSHEA *et al.* 2013). MIG-15 (NIK kinase) is a cytoplasmic protein and evidence indicates that *mig-15* functions cell-autonomously to mediate a response to UNC-6 (POINAT *et al.* 2002; TEULIÈRE *et al.* 2011). It’s proposed that *mig-15* acts with *unc-5* to polarize the growth cone’s response and that it controls the asymmetric localization of MIG-10 and UNC-40 (TEULIÈRE *et al.* 2011; YANG *et al.* 2014). We previously noted that HSN neurons often become bipolar in *mig-15* mutants and frequently UNC-40::GFP is localized to multiple surfaces in a single neuron, suggesting that loss of MIG-15 enhances the ability of UNC-40::GFP to cluster (YANG *et al.* 2014). In our experiments we used the *mig-10 (ct141)* loss-of-function allele (MANSER AND WOOD 1990; MANSER *et al.* 1997) and the *mig-15(rh148)* allele, which causes a missense mutation in the ATP-binding pocket of the kinase domain and is a weak allele of *mig-15* (SHAKIR *et al.* 2006; CHAPMAN *et al.* 2008).

We find that the extra processes induced by UNC-6ΔC expression are suppressed by *mig 10(ct141)* (Figurs 7F). We also find that the *mig-15* mutation causes extra HSN processes and that the loss of UNC-5 function suppresses these extra HSN processes (Figures 7F and 7G). These results support the hypothesis that the ability of UNC-5 to regulate the development of multiple protrusions involves the molecular machinery that controls UNC-40-mediated asymmetric neuronal outgrowth.

### UNC-5 is required for PLM overextension

The SDAL model predicts that the ability of UNC-5 to regulate the length and number of neural protrusions is independent of the direction of outgrowth. HSN sends a single axon ventrally, while PLM sends an axon anteriorly from a posteriorly positioned cell body. The HSN axon travels towards UNC-6 sources, whereas the PLM axon pathway is perpendicular to UNC-6 sources. To investigate whether UNC-5 activity can regulate the length or number of processes that develop perpendicular to UNC-6 sources we examined the development of the PLM axon. We also chose PLM because UNC-5 was already known to affect the length of the PLM axon (LI *et al.* 2008).

Given that UNC-5 activity is involved in the overextension of the PLM axon, and that the *mig-15* mutation affects HSN outgrowth in an UNC-40 dependent fashion, we decided to test whether PLM overextension might be induced by the *mig-15* mutation in an UNC-40-dependent fashion. The HSN results suggest that altering *mig-15* function creates a sensitized genetic background. That is, the *unc-5(ev480)* mutation suppresses HSN outgrowth extension in both the wild-type and *mig-15(rh148)* backgrounds, but the *mig-15* mutation creates a stronger patterning phenotype. This idea is supported by the evidence that the *mig-15* mutation enhances the ability of UNC-40 to localize at surfaces (YANG *et al.* 2014).

We find that in *mig-15(rh148)* mutants the PLM axon often fails to terminate at its normal position and instead extends beyond the ALM cell body. This overextension is suppressed in *unc-5(e53);mig-15(rh148)* and *unc-40(e1430);mig-15(rh148)* mutants (Figures 8A and 8B). The results are consistent with the idea that UNC-5 is required for the UNC-40-mediated outgrowth activity that causes overextension in *mig-15(rh148)* mutants.

**Figure 8.**
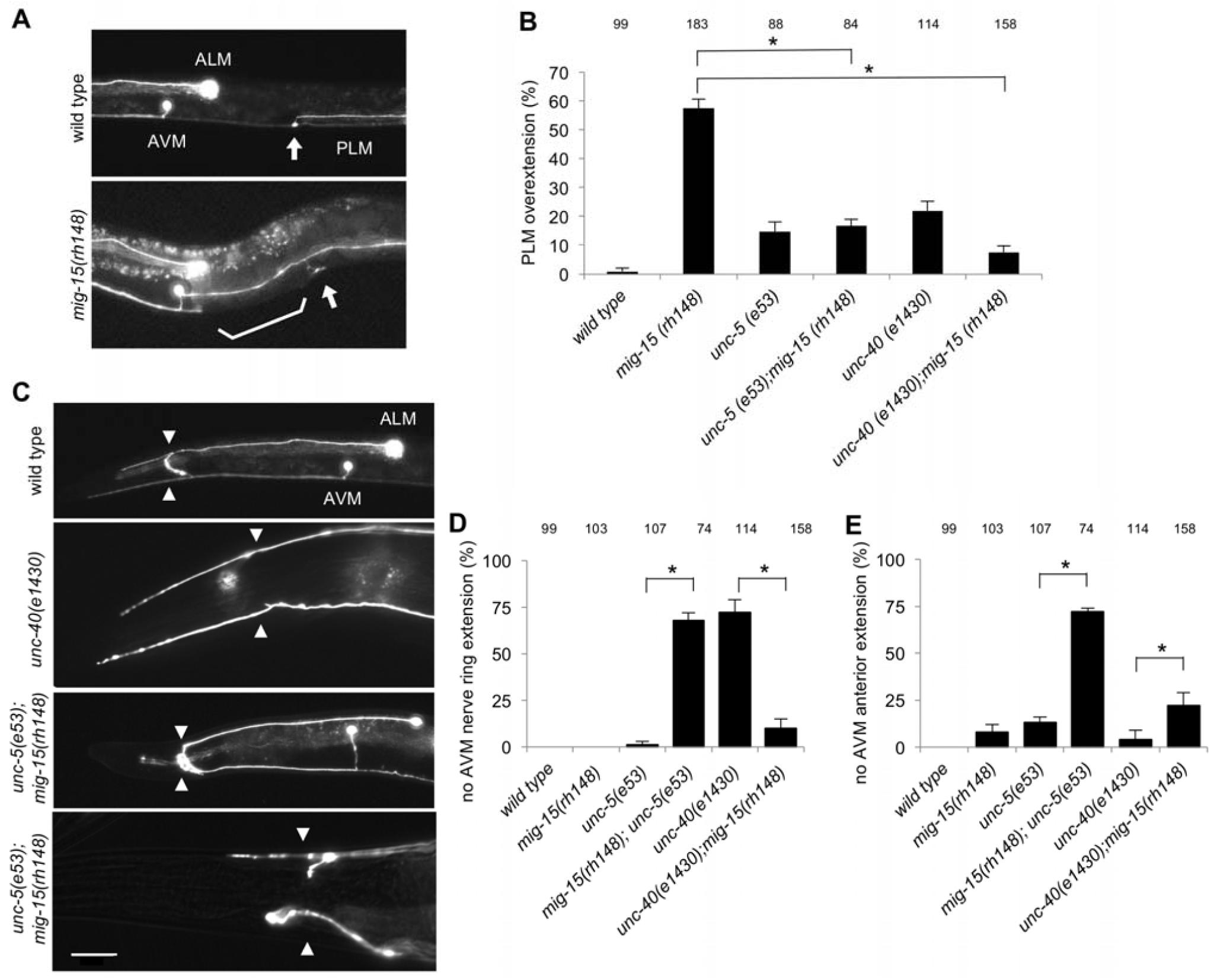
UNC-5 regulates the patterning of extension from ALM, AVM, and PLM. **(A)** Photomicrographs of the ALM, AVM, and PLM neurons at the L4 stage in wild-type animals and *mig-15* mutants. In wildtype (top) a single PLM axon travels anteriorly from the posterior cell body (not shown). Near the vulva (arrow) the axon branches; one branch extends to the ventral nerve chord and another extends anteriorly. The anterior extension terminates before reaching the area of the ALM cell body. In *mig-15* mutants the PLM can extend anteriorly past the ALM cell body (bottom). **(B)** The percentage of PLM neurons where the PLM neuron extend anteriorly past the ALM cell body. The anterior extension often over-extends in *mig-15* mutants. Loss of *unc-5* or *unc-40* function can suppress this phenotype. **(C)** Photomicrographs of the ALM and AVM neurons at the L4 stage in wild-type animals and mutants showing different patterns of outgrowth extension. In wildtype (top) a single axon travels anteriorly to the nerve ring (arrowheads). At the nerve ring the axon branches; one branch extends further anteriorly and the other extends into the nerve ring. In mutants, one or both axons may only extend anteriorly and will not extend into the nerve ring (second from top). Or one or both axons will only extend into the nerve ring and will not extend anteriorly (third from top). Or one or both axons will fail to extend into either the nerve ring or anteriorly (bottom). Scale bar: 20 μm. **(D)** The percentage of AVM neurons where the AVM neuron failed to extend into the nerve ring. The neuron often fails to extend in the *unc-40* and *mig-15;unc-5* mutants, whereas it does extend in the *mig-15, unc 5,* and *mig-15;unc-40* mutants. Error bars indicated the standard error mean; n values are indicated above each column. Significant differences (two-tailed t-test), *P<0.001. **(E)** The percentage of AVM neurons where the AVM neuron failed to extend anteriorly, past the nerve ring. The neuron often fails to extend anteriorly in the *mig-15;unc-5* mutants, whereas it does extend in the *mig-15, unc-5, unc-40,* and *unc-40;mig-15* mutants. There is a significant difference between the *unc-40* and *unc-40;mig-15* mutants.

**Figure 9.**
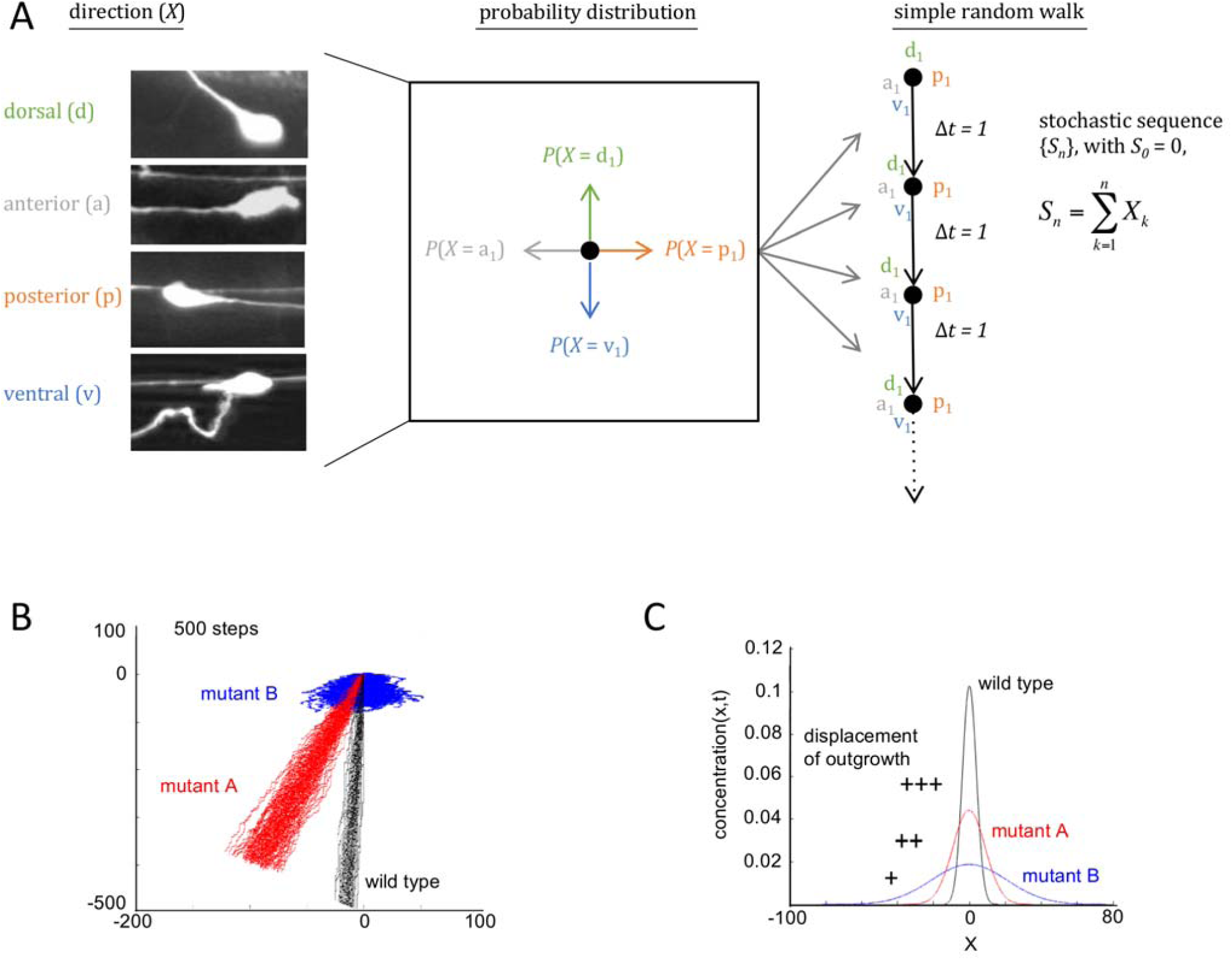
Assay to measure the effects a mutation has on movement. (**A**) The direction of outgrowth extension from the HSN cell body can vary and whether the axon developed in the dorsal, anterior, posterior, or ventral direction in L4 stage animals is scored (left panel). This creates a probability distribution in which the direction (*X*) is a random variable (center panel). A simple random walk is generated by using the same probability distribution for a succession of steps with an equal time interval (right panel). (**B**) For wildtype and two mutants, 50 simulated random walks of 500 steps were plotted from an origin (0,0). The results graphically indicate the directional bias for movement. For random walk movement created in mutant A (red, results from *unc-5(e53)*), the directional bias is shifted anteriorly (left) relative to wildtype. The results also graphically show the displacement of movement. For random walk movement created in mutant B (blue, results from *egl-20(n585);sax-3(ky123)*), the average of the final position (displacement) from the origin is a much shorter distance than wildtype. (**C**) Plots of the normal distribution of the final position along the x axis of the random walk tracks shown in B. The mean position for each is set at 0. The plots graphically illustrate how random walks constructed from the probability distribution for the direction of outgrowth extensions can reveal a diffusion process.

### UNC-5 is required for ALM and AVM branching and extension

We also investigated the effect of UNC-5 activity on patterning where sources of UNC-6 and other cues are in a more complex arrangement. Specifically, we examined whether UNC-5 plays a role in the outgrowth of AVM and ALM processes at the nerve ring. During larval development, processes from the AVM neuron and the two ALM neurons (one on each side of the animal) migrate anteriorly to the nerve ring at dorsal and ventral positions respectively (Figure 8C). At the nerve ring each axon branches; one branch extends further anteriorly and the other extends into the nerve ring. Evidence suggests that at the midbody of the animal the positioning of these axons along the dorsal-ventral axis requires UNC-6, UNC-40, and UNC-5 activity. In *unc-6*, *unc-40,* and *unc-5* null mutants, or when the UNC-6 expression pattern is altered, the longitudinal nerves are mispositioned (REN *et al.* 1999). Glia cells and neurons at the nerve ring are sources of UNC-6 (WADSWORTH *et al.* 1996). The guidance of some axons in the nerve ring are disrupted in *unc-6* and *unc-40* mutants (HAO *et al.* 2001; YOSHIMURA *et al.* 2008). The precise spatial and temporal arrangement of the UNC-6 cue in relationship to the position of the migrating growth cones is not fully understood. Nevertheless, the anteriorly migrating growth cones appear to use the UNC-6 cue from the ventral sources to help maintain the correct dorsal-ventral position, even while moving towards the nerve ring, which is a new source of UNC-6 that is perpendicular to the ventral source. At the nerve ring the axons branch. One process continues anteriorly, moving past the new UNC-6 source, whereas the other projects at a right angle and moves parallel to the new source.

We find genetic interactions involving *unc-5, unc-40,* and *mig-15* that affect outgrowth patterning of the ALM and AVM extensions at the nerve ring (Figures 8C, 8D, and 8E). In *mig 15(rh148);unc-5(e53)* mutants, the AVM axon often fails to extend anteriorly from the branch point and only extends into the nerve ring, or it fails to extend into the nerve ring and only extends anteriorly, or it fails to do both and terminates at this point. In *unc-40(e1430)* mutants, the axon often fails to branch into the nerve ring, although it extends anteriorly. In comparison, in *unc-40(e1430);mig-15(rh148)* mutants more axons extend into the nerve ring. These results suggest that UNC-5 (and MIG-15) helps regulate UNC-40-mediated outgrowth to pattern the outgrowth at the nerve ring.

### Interactions between *unc-5* and other genes affect a probability distribution for the direction of extension

We hypothesize that there are interactions between *unc-5* and other genes that control the degree to which the direction of outgrowth fluctuates. Probability distributions for the direction of extension are used to study how genes affect the fluctuation of outgrowth activity. By comparing the distributions created from wild-type and mutant animals, the relative effect that genes have on the fluctuation can be determined. To accomplish this, the direction that the HSN axon initially extends from the cell body is scored (Figure 9A).

Using this assay, we examined genetic interactions between *unc-5* and four other genes; *elg-20, sax-3, madd-2,* or *unc-6.* We have chosen these particular genes because previous observations suggest interactions. 1) EGL-20 (Wnt) is a secreted cue expressed from posterior sources (PAN *et al.* 2006) and it affects to which surface of the HSN neuron the UNC-40 receptor localizes and mediates outgrowth (KULKARNI *et al.* 2013). Based on a directional phenotype, a synergistic interaction between *unc-5* and *egl-20* has been observed. In either *unc-5* or *egl-20* mutants the ventral extension of AVM and PVM axons is only slightly impaired, whereas in the double mutants there is much greater penetrance (LEVY-STRUMPF AND CULOTTI 2014). 2) SAX-3(Robo) is a receptor that regulates axon guidance and is required for the asymmetric localization of UNC 40 in HSN (TANG AND WADSWORTH 2014). Based on a directional phenotype, SAX-3 and UNC-40 appear to act in parallel to guide the HSN towards the ventral nerve cord (XU *et al.* 2015). 3) MADD-2 is a cytoplasmic protein of the tripartite motif (TRIM) family that potentiates UNC-40 activity in response to UNC-6 (ALEXANDER *et al.* 2009; ALEXANDER *et al.* 2010; HAO *et al.* 2010; MORIKAWA *et al.* 2011; SONG *et al.* 2011; WANG *et al.* 2014). MADD-2::GFP and F-actin colocalize with UNC-40::GFP clusters in the anchor cell (WANG *et al.* 2014). 4) Of course, UNC-6 is an UNC 5 ligand. DCC (UNC-40) and UNC5 (UNC-5) are thought to act independently or in a complex to mediate responses to netrin (UNC-6) (COLAVITA AND CULOTTI 1998; HONG *et al.* 1999; MACNEIL *et al.* 2009; LAI WING SUN *et al.* 2011).

In a test for interaction with *egl-20*, we find that in comparison to *unc-5(e53)* or *egl-20(n585)* mutants, the *unc-5(e53);egl-20(n585)* double mutant have a lower probability for ventral outgrowth and higher probability for outgrowth in other directions (Table 1). This suggests that *unc-5* and *egl-20* may act in parallel to achieve the highest probability for HSN ventral outgrowth, *i.e.* they act to prevent UNC-40-mediated outgrowth from fluctuating in other directions.

In a test for interaction with *sax-3,* we find that the probability of outgrowth in each direction in *unc-5(e53);sax-3(ky200)* mutants is similar to the probabilities in *sax-3(ky200)*or *sax-3(ky123)* mutants (Table 1). Given the results with *unc-5* and *egl-20*, we further tested the probability of outgrowth in each direction in *egl-20(n585);sax-3(ky123)* mutants. We find that it is similar to the probabilities in *sax-3(ky200)*or *sax-3(ky123)* mutants (Table 1). The *sax-3(ky123)* allele results in a deletion of the signal sequence and first exon of the gene, whereas sax-3(ky200) contains a missense mutation which is thought to cause protein misfolding and mislocalization at the restrictive temperature (25°C) (ZALLEN *et al.* 1998; WANG *et al.* 2013). The *egl 20(n585);sax-3(ky123)* mutants do not grow well and so it is easier to use the temperature sensitive *sax-3* allele. Together, the results suggest that *sax-3* may be required for both the *unc-5-* and the *egl-20-*mediated activities that allow the highest probability for HSN ventral outgrowth.

In a test for interaction with *madd-2,* we find that the probability of outgrowth in each direction in *unc-5(e53);madd-2(tr103)* mutants is similar to the probabilities in *madd-2(tr103)* mutants (Table 1). There is a higher probability for anterior HSN outgrowth, similar to what is observed in *unc 40(e1430)* mutants. These results suggest that *madd-2* might be required for the *unc-40* outgrowth activity. The probability of outgrowth in each direction in *madd-2(tr103);sax 3(ky123)* mutants is similar to the probabilities in *sax-3(ky200)*or *sax-3(ky123)* mutants (Table 1). The *madd-2(tr103)* allele appears to act as a genetic null (ALEXANDER *et al.* 2010).

In a test for interaction with *unc-6,* we find that the probability of outgrowth in each direction in *unc-5(e53);unc-6(ev400)* and *unc-40(e1430);unc-5(e53)* mutants is similar to the probabilities in *unc-6(ev400)* mutants insofar as there is a lower probability for ventral outgrowth and a higher probability for anterior outgrowth (Table 1). However, the probabilities in each direction are closer to those obtained from the *unc-40(e1430)* mutants because the probability of anterior outgrowth is lower in these mutants than in *unc-6* mutants. This suggest that UNC-5 and UNC-40 might help increase the probability of anterior outgrowth in the absence of UNC-6.

### *unc-5* is a member of a class of genes that has a similar effect on the spatial extent of movement

The results above show that *unc-5* and its interactions with other genes affect the degree to which the direction of outgrowth fluctuates. The degree of fluctuation differs depending on the genes involved. A property of random movement is that the more the direction of movement fluctuates, the shorter the distance of travel is in a given amount of time (Figure 2E). To depict how *unc-5* and other genes differentially regulate the spatial extent of movement, we use random walk modeling. Random walks describe movement that occurs as a series of steps in which the angles and the distances between each step is decided according to some probability distribution. By using the probability distribution obtained from a mutant for each step of a random walk, and by keeping the distance of each step equal, a random walk can be constructed (Figure 9A). In effect, this method applies the probability distribution to discrete particles having idealized random walk movement on a lattice. By plotting random walks derived from wild-type animals and different mutants, the relative effect that mutations have on random walk movement can be visualized. For example, Figure 9B shows 50 tracks of 500 steps for wildtype and two mutants (mutant A is *unc-5(e53)* and mutant B is *egl-20(n585);sax 3(ky123)*). This reveals the effect that a mutation has on the displacement of movement. After 500 steps the displacement from the origin (0,0) is on average less for mutant A than for wildtype, and less for mutant B than for wildtype or mutant A.

The random walk models show the relative effect that a mutation has on a property of outgrowth movement. It is worth noting that this is not modeling the actual trajectory of migrating axons. As discussed in the introduction, neuronal outgrowth is essentially a mass transport process in which mass (the molecular species of the membrane) is sustained at the leading edge and moves outward. Our assay compares the effect that different mutations would have on the movement of mass at the leading edge of an extension if the conditions of the system were kept constant. Of course, *in vivo* the conditions are not constant. For one, as an extension moves it will encounter new environments where the cues may be new or at different concentrations, all of which affect the probability distribution. The actual patterns of outgrowth observed are the result of all the probabilities for outgrowth that occur at each instance of time. It has recently been suggested that our description might be more accurately described as neuro-percolation, a superposition of random-walks (AIELLO 2016).

Our random walk analysis compares the effect that different mutations have on the properties of movement. In wild-type animals, there is a high probability for outgrowth in the ventral direction. The analysis shows that conditions in wildtype create nearly straight-line movement, *i.e.* if the same random walk is repeatedly done for the same number of steps, starting at the same origin, the final position of the walk along the x axis does not vary a great amount. In comparison, we find that a mutation can create random walk movement in which the final position is more varied. This variation occurs because the mutation increases the probability of outgrowth in other directions. For each mutation, we simulate 50 random walks of 500 steps and derive the mean and standard deviation of the final position along the X-axis. To compare strains, we plot the normal distribution, setting the mean at the same value for each. The difference between the curve for a mutant and wildtype shows the degree to which the mutation caused the direction of outgrowth to fluctuate (Figure 9C).

The results reveal four different distribution patterns (Figure 10). The first class is the wild-type distribution, which has the distribution curve with the highest peak. The second class comprises *unc-5, egl-20, unc-53,* and *unc-6* in which the distribution curve is flatter than the wild-type curve. We included *unc-53* because our previous study showed that it has genetic interactions with *unc-5* and *unc-6* (KULKARNI *et al.* 2013). The *unc-53* gene encodes a cytoskeletal regulator related to the mammalian NAV proteins and *unc-53* mutations cause guidance defects (MAES *et al.* 2002; STRINGHAM *et al.* 2002; STRINGHAM AND SCHMIDT 2009). The third class has a distribution curve which is flatter than the second and comprises *sax-3, mig-15,* and several double mutation combinations (Figure 10). The fourth class has the flattest distribution curve and comprises *egl-20;sax-3*, *unc-40;sax-3,* and *unc-53;sax-3;unc-6*. This class indicates the greatest degree of fluctuation. The ability to cause the direction of movement to fluctuate is not associated with a specific direction of HSN movement. For example, *unc-5;sax-3, unc-53;unc-6, unc-40;egl-20,* and *madd-2;sax-3* each show a widely dispersed pattern, but the direction is ventral, dorsal, anterior, and posterior, respectively (Figure 10).

**Figure 10.**
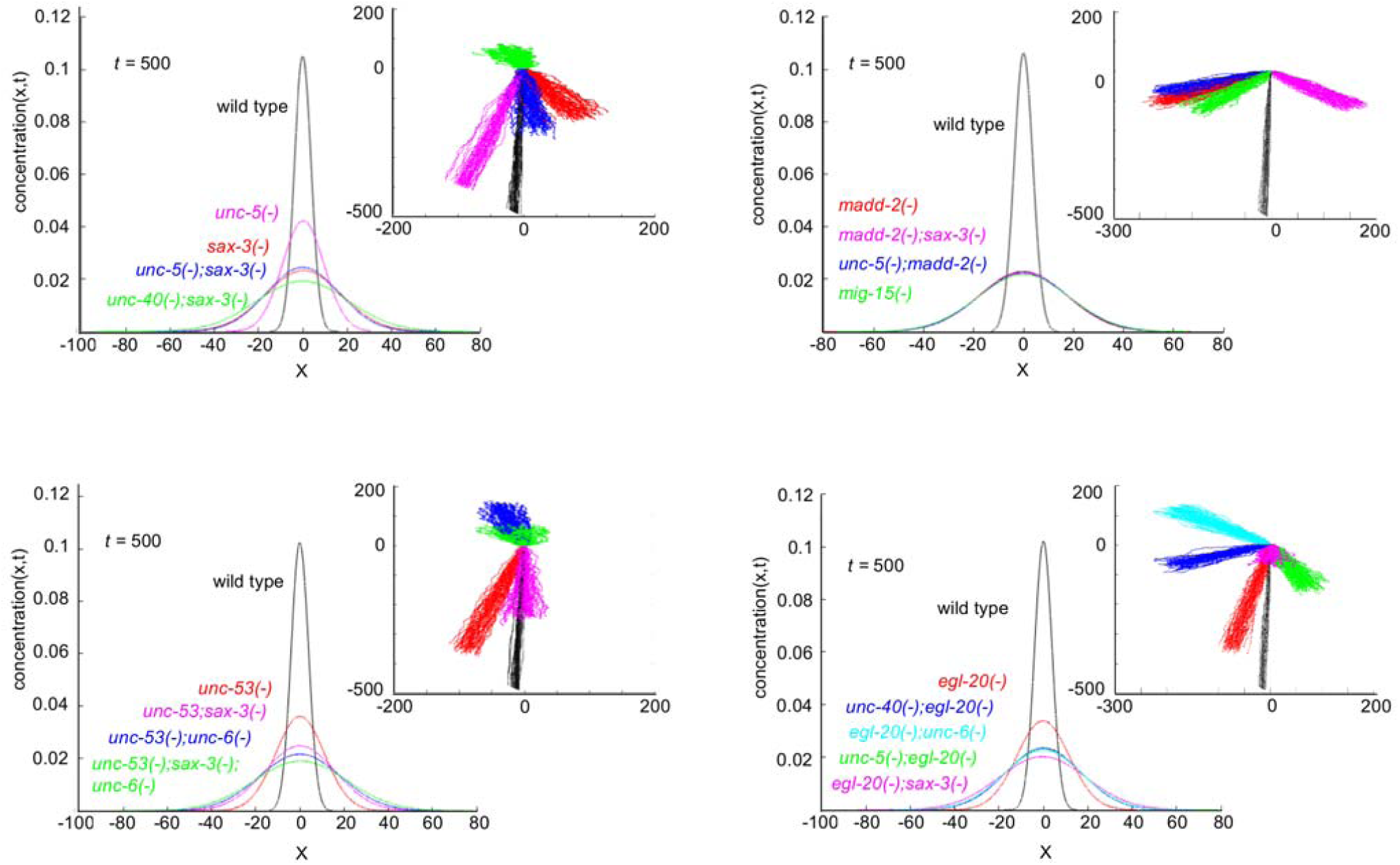
Mutations have different effects on movement. Examples of random walk analyses using the direction of axon development from the HSN neuron in different mutants (Table 1). The graphs were created as described in the figure legend of Figure 9. For each panel, plots are shown for the normal distribution of the final position along the x axis for the random walk tracks plotted in the inserts. The inserts depict the random walk movement that would be produced by the probability distribution for the direction of outgrowth in the mutant. Plots derived from the same data are colored alike. Each panel depicts the analyses of four different mutants and wildtype. Three different distribution patterns are observed: (1) the wild-type distribution, which has the distribution curve with the highest peak; (2) the *unc-5, egl-20, unc-53,* and *unc-6* (not shown) distribution, which is flatter than the wild-type curve; (3) the *madd-2, sax-3, mig-15,* and double combinations, which have the flattest distribution curve.

The distribution patterns indicate that genes have different effects on the extent that outgrowth movement can travel through the environment. Mean squared displacement (MSD) is a measure of the spatial extent of random motion. The MSD can be calculated from the random walk data. Plotting MSD as a function of the time interval shows how much an object displaces, on average, in a given interval of time, squared (Figure 11A). For normal molecular diffusion, the slope of the MSD curve is directly related to the diffusion coefficient. In cell migration models this value is referred to as the random motility coefficient. Coefficients are experimentally determined; they describe how long it takes a particular substance to move through a particular medium. We determine this value in order to numerically and graphically compare how mutations can alter displacement relative to wildtype (Figure 11B). The four classes of genes are apparent by comparing the height of the bars in Figure 11B. Results for the *unc-40* mutation are also show. The random walk pattern is published (TANG AND WADSWORTH 2014).

**Figure 11.**
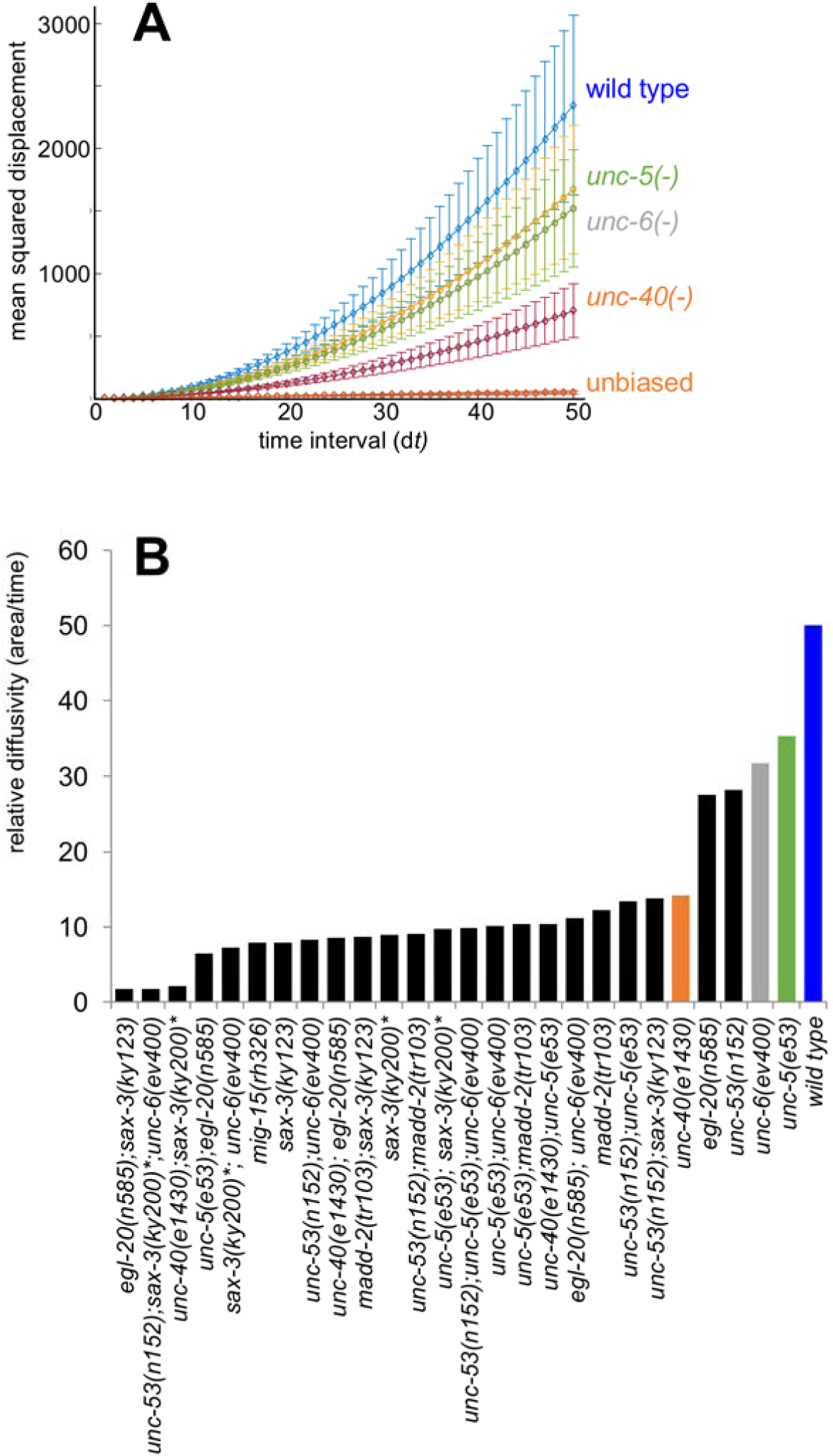
Mutations alter the spatial extent of movement. **(A)** Plotted are the mean squared displacement (MSD) curves as a function of time interval (d*t*). The values are in arbitrary units, since the time scale was arbitrarily set at 1. The curves show the extent that different mutations can alter the MSD relative to wildtype and the MSD caused by an unbiased random walk. For each time interval, mean and s.e.m. are plotted. **(B)** From the slope of MSD curves a coefficient can be derived that gives the relative rate of diffusion. Colored bars correspond to the like-colored curves given in panel A. The coefficients for *unc-5, egl-20, unc-53,* and *unc-6* form a class that is distinct from that derived from wildtype and from the double mutants.

The results of this modeling suggest that the activities of certain genes, and combinations of genes, have distinct effects on the rate of outgrowth movement. In theory, these differences could be an important means by which genes cause different outgrowth patterns.

### UNC-40 receptor clustering is coupled to the SDAL process

We investigated the relationship between UNC-40::GFP localization and outgrowth movement. Beginning in the early L2 stage, UNC-40::GFP becomes localized to the ventral side of HSN in wildtype (ADLER *et al.* 2006; KULKARNI *et al.* 2013). Reflecting the dynamic morphological changes that occur as the HSN axon forms, the site of asymmetric UNC-40::GFP localization alternates in the neurites and along the ventral surface of the neuron (KULKARNI *et al.* 2013). Dynamic UNC-40::GFP localization patterns have also been reported for the anchor cell, in which UNC-40 and UNC-6 are also key regulators of extension (ZIEL *et al.* 2009; HAGEDORN *et al.* 2013). Live imaging of the anchor cell reveals that UNC-40::GFP “clusters” form, disassemble, and reform along the membrane (WANG *et al.* 2014). However, live imaging can’t directly ascertain whether the position of a cluster is randomly determined since a movement event cannot be repeatedly observed to determine a probability distribution. Mathematical modeling of cluster movement as a stochastic process has not been done.

The UNC-40::GFP clustering phenomena raises questions about the relationship between robust UNC-40 clustering (*i.e.,* sites of distinct UNC-40 localization observable by UNC-40::GFP) and UNC-40-mediated outgrowth activity. Two models are presented in Figure 12. In the first model, the output of the SDAL process is receptor clustering (Figure 12A). After a cluster becomes stabilized at a site, the machinery required for outgrowth is recruited and outgrowth occurs. In our model, the SDAL process and UNC-40-mediated outgrowth activity are coupled and are part of the same stochastic process that occurs at the micro-scale (Figure 12B). UNC 40::GFP clustering is a macro-scale event which can be observed. It is a consequence of the micro-scale events.

**Figure 12.**
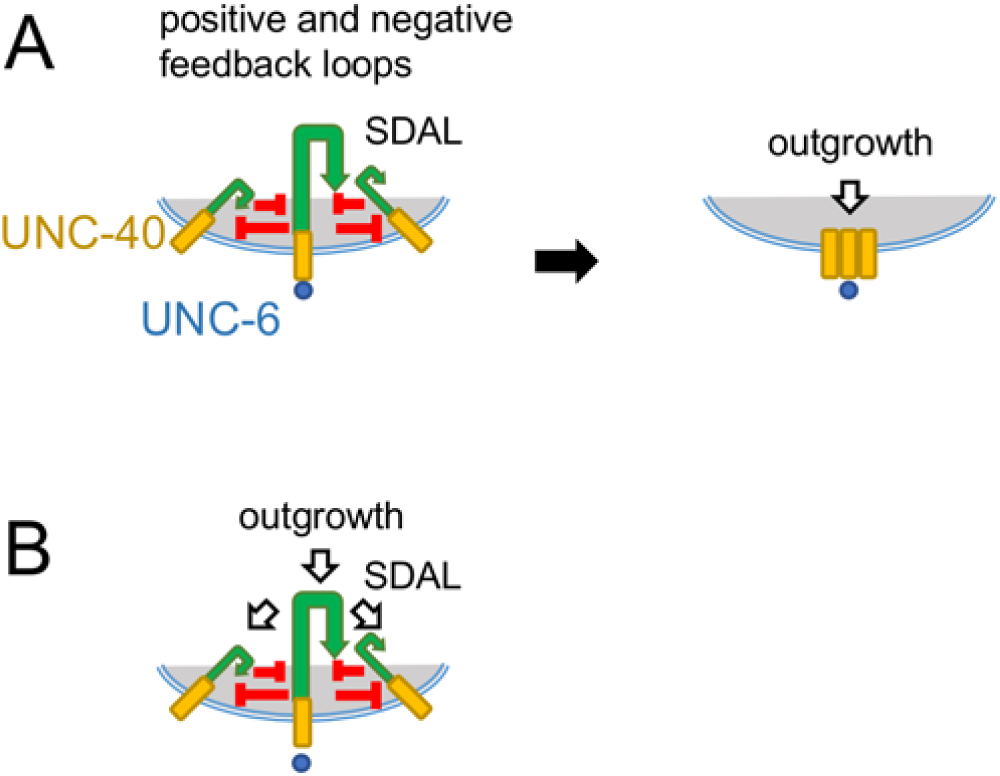
Models for the relationship between UNC-40-mediated outgrowth activity and UNC-40 receptor clustering. **(A)** The SDAL process is illustrated as in Figure 1. In this model, the self-organizing UNC-40 SDAL process causes observable UNC-40 receptor clustering. UNC-6 stabilizes receptor clustering at a site and the outgrowth machinery is then recruited to cause outgrowth at the site. Although the initial direction of asymmetric receptor localization is determined stochastically, the direction of outgrowth is determined by the site of stabilization. **(B)** In this model, the self-organizing UNC-40 SDAL process is coupled to the outgrowth machinery. The direction of both asymmetric receptor localization and outgrowth activity are stochastically determined. Observable receptor clustering arises as the result of the process because receptor localization can become successively concentrated to a smaller area over time. Cluster formation is an observable phenomenon of the process, not a prerequisite for outgrowth activity. This model postulates innumerable fluctuating sites that generate force in various directions along the membrane.

The models make specific predictions that can be tested. In the first model, UNC-40-mediated outgrowth will not happen if UNC-40 does not cluster. In our model, the loss of UNC-40 clustering does not lead to a loss of UNC-40-mediated outgrowth. In the *sax-3* mutant there is a large fluctuation in the direction of outgrowth; it is in the third class of mutants (Figures 10 and 11). We previously reported that *sax-3* is required for robust UNC-40::GFP asymmetric localization; in *sax-3* mutants UNC-40::GFP remains uniformly dispersed around the periphery of HSN ((TANG AND WADSWORTH 2014) and Figure 13). Whereas in the *sax-3* mutant there is a ventral bias for outgrowth, in the *unc-40;sax-3* mutant there is not (Figure 10). This suggests that in the *sax-3* mutant there is UNC-40-mediated outgrowth activity that helps create a ventral bias. This is consistent with our model because UNC-40-mediated outgrowth activity is occurring even when robust UNC-40::GFP is not observed.

**Figure 13.**
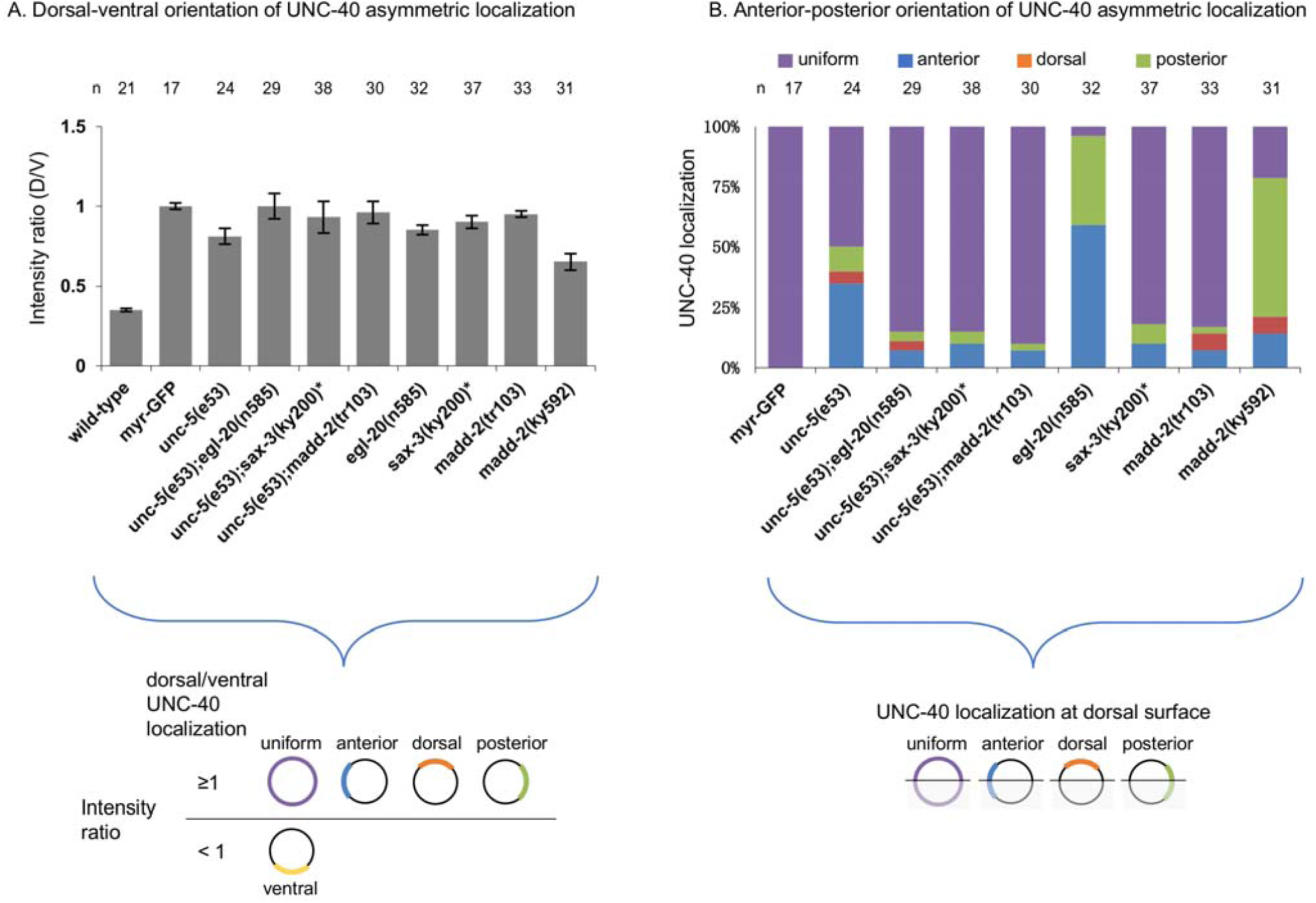
Mutations affect asymmetric intracellular UNC-40::GFP localization. **(A)** Graph indicating the dorsal-ventral localization of UNC-40::GFP in HSN. The graph shows the average ratio of dorsal-to-ventral intensity from linescan intensity plots of the UNC-40::GFP signal around the periphery of the HSN cell. UNC-40::GFP is ventrally localized in wildtype, but the ratio is different in the mutants. Error bars represent standard error of mean. Below is a graphic representation of the possible UNC-40 localization patterns when the intensity ratio is ≥ 1 or is <1. **(B)** Graph indicating the anterior-posterior localization of UNC-40::GFP. To determine orientation, line-scan intensity plots of the UNC-40::GFP signal across the dorsal periphery of the HSN cell were taken, the dorsal surface was geometrically divided into three equal segments, and the total intensity of each was recorded. The percent intensity was calculated for each segment and ANOVA was used to determine if there is a significant difference between the three segments (see Material and Methods). The graph shows the percent of animals that had significant localization in the indicated segments or that had uniform distribution. Whereas in the *unc-5* and *egl-20* mutants there is a bias for anterior or posterior localization, there is a uniform distribution in *unc-5;egl-20* double mutants. Uniform distribution is also observed in strong loss-of function *sax-3* and *madd-2* mutants. (*) Animals grown at the *sax-3(ky200)* restrictive temperature (25°C). Below each graph is a representation of the possible UNC-40 localization patterns. The orientation of UNC-40 localization is color-coded as in B.

We hypothesize that a consequence of the micro-scale SDAL process over time is macro-scale UNC-40 clustering. If so, then *unc-5* activity should affect UNC-40::GFP clustering because it affects the degree to which the direction of UNC-40 receptor localization fluctuates. However, even though there is a higher probability that localization occurs at surfaces other than at the ventral surface, we observe robust asymmetrically localized UNC-40::GFP clustering in *unc 5(e53)* mutants (KULKARNI *et al.* 2013). We speculate that *unc-5(e53),* as well as other gene mutations, do not cause the direction of UNC-40 localization to fluctuate enough to prevent observable UNC-40::GFP clustering. We therefore decided to examine UNC-40::GFP clustering in double mutants to determine whether the ability to observe UNC-40::GFP clustering is correlated with the degree of fluctuation.

We made double mutant combinations between *unc-5*, and *egl-20* or *unc-53.* In *egl-20* and *unc 53* single mutants there is fluctuation in the direction of outgrowth (Figures 10 and 11) and robust asymmetrical UNC-40::GFP localization (Figure 13, *unc-53* results were previously reported (KULKARNI *et al.* 2013)). In comparison to the single mutants, the double mutants all show an increase in the degree to which the direction of outgrowth fluctuates (Figures 10 and 11). Further, in contrast to the single mutants, UNC-40::GFP remains uniformly dispersed around the periphery of HSN in the double mutants (Figure 13). The results suggest a correlation between increased fluctuation of UNC-40-mediated outgrowth activity and the ability to detect UNC-40::GFP clustering. This is consistent with our model (Figure 12B). We also observe that in *madd-2(tr103)* mutants the direction of outgrowth fluctuates (Table 1), but unlike *egl-20* and *unc-53* single mutants, there is not robust asymmetrical UNC-40::GFP localization and UNC-40::GFP remains uniformly dispersed (Figure 13). The double mutants, *unc-5;madd-2,* are similar to the single *madd-2* mutant. Similar results are observed with *sax-3* and *unc-5;sax-3* mutants (Figure 13). We hypothesize that in the *madd-2* and *sax-3* mutations the degree to which the direction of UNC-40 localization fluctuates is so great that the *unc-5* mutation makes no difference on the UNC-40::GFP clustering phenotype.

## Discussion

We have proposed a model of neuronal outgrowth movement that is based on statistically dependent asymmetric localization (SDAL). This model states that the probability of UNC-4O localizing and mediating outgrowth at one site affects the probability of localization and outgrowth at other sites as well. By regulating this process, genes control the degree to which the direction of outgrowth fluctuates and, consequently, the outward movement of the plasma membrane. UNC-5 is a receptor for UNC-6 and can form a complex with UNC-40. UNC-5 is commonly proposed to direct outgrowth by mediating a repulsive response to UNC-6. In contrast, our model is not based on the concept of repulsion and it predicts that UNC-5 can control the rate of outward movement that is directed towards, away from, or perpendicular to UNC-6 sources. We report that *unc-5* loss-of-function mutations affect the development of multiple neurites that develop from HSN and extend towards UNC-6 sources. They also suppress the development of extra HSN processes which are induced by a *mig-15* mutation or by expression of the N-terminal fragment of UNC-6 and which extend towards UNC-6 sources. We also observe that *unc-5* mutations suppress the anterior overextension of the PLM axon that occurs in the *mig-15* mutant. This axon extends perpendicular to UNC-6 sources. Finally, *unc-5* loss-of-function mutations affect the branching and extension of ALM and AVM axons at the nerve ring where the sources of UNC-6 are in a more complex arrangement. Below we discuss how the SDAL model can be used to interpret *unc-5* mutant phenotypes. We argue that in each case, phenotypes can be explained by the ability of UNC-5 to affect UNC-40 asymmetric localization, which in turn controls the degree to which the direction of outgrowth activity fluctuates and the extent of outward movement. Our model also suggests genes that were previously classified as regulating attraction or repulsion might act with *unc-5* to regulate neuronal outgrowth by controlling the degree to which the direction of UNC-40-mediated outgrowth fluctuates. We show that UNC-5 acts together with the cytoplasmic protein UNC-53 to regulate UNC-40 asymmetric localization in response to the UNC-6 and EGL-20 extracellular cues.

**PLM extension phenotype:** We hypothesize that cue(s) present around the PLM cell body create a strong bias for anterior outgrowth activity (Figure 14A). These include UNC-6 and other cues that flank the longitudinal pathway and cause an equal probability of outgrowth in the dorsal and ventral directions. UNC-40 SDAL activity acts to suppress nonUNC-40 SDAL activity (Figure 6). As the extension moves towards more anterior positions (Figure 14A, positions 2 and 3), it encounters higher levels of a cue(s) that promotes outgrowth through nonUNC-40 receptors. As a result, the probability of nonUNC-40 SDAL activity at the dorsal and ventral surfaces of the leading edge increases. Because the asymmetric localization of a receptor is statistically dependent, the probability of nonUNC-40 SDAL activity at the anterior surface of the leading edge must decrease as the localization elsewhere increases. While this effect does not necessarily change the anterior bias for outgrowth, it does significantly increases the degree to which the direction of nonUNC-40 outgrowth activity fluctuates, which consequently decreases the extent of outward movement (Figure 14C). This effect stalls forward movement.

**Figure 14.**
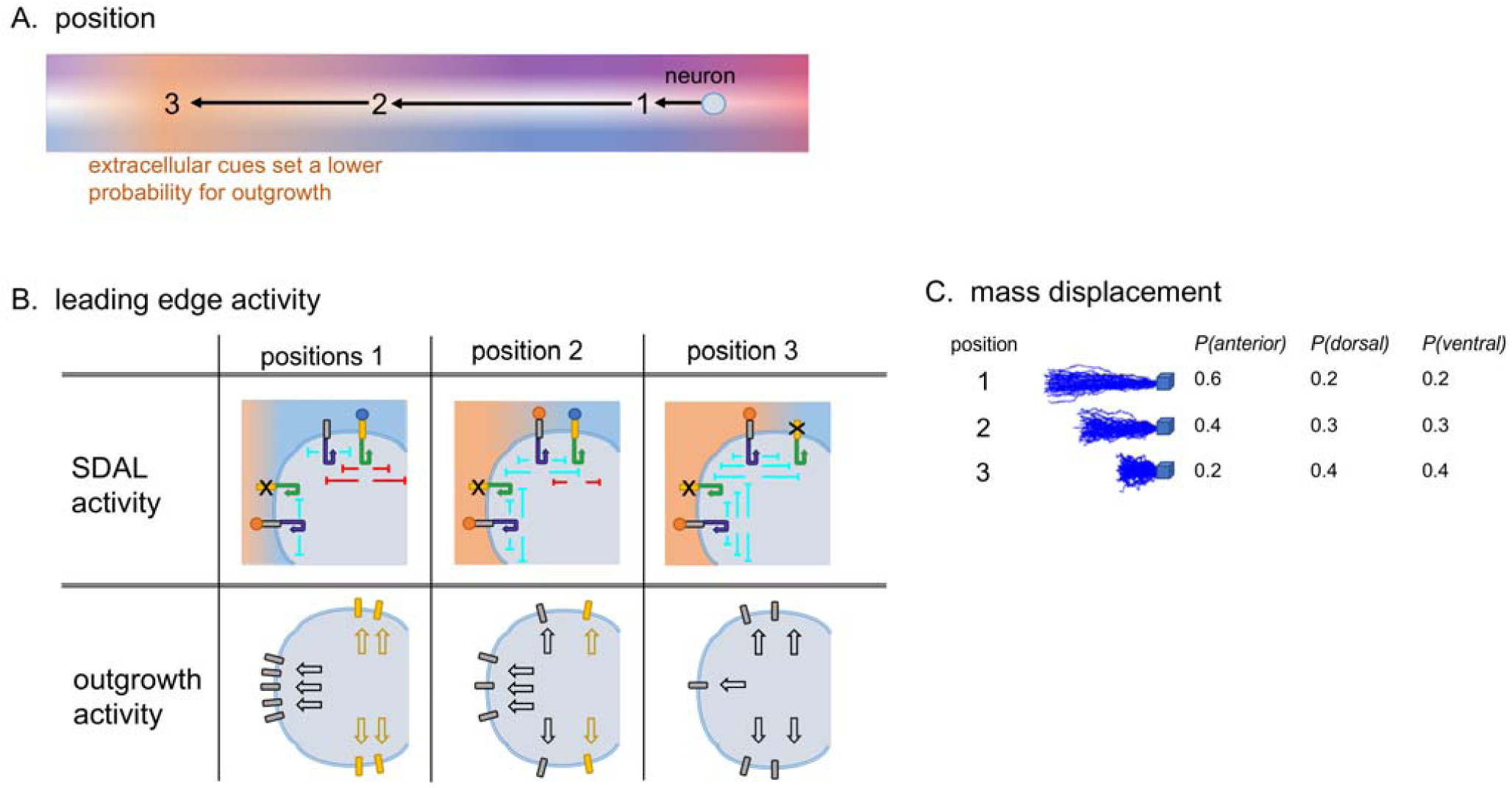
Model for the outgrowth movement of PLM. Schematic diagrams of the anteriorly directed outgrowth of PLM. The features of the schematic are presented in Figure 3. **(A)** An extension encounters different levels of extracellular cues at each of three positions (1-3). Cues dorsal and ventral of the pathway maintain an equal probability of outgrowth in both directions. The extension encounters increasing levels of an extracellular cue(s) as it moves towards position 3. At position 3, a cue(s) prevent further anterior extension in wildtype. **(B)** Table showing for each of the positions depicted in A the positive feedback (arrows) and negative feedback (lines) associated with SDAL activity (see Figure 6) and the predicted effect the SDAL activity has on outgrowth activity. At position 1, strong UNC-40 activity along the dorsal and ventral (not shown) facing surfaces of the leading edge inhibit nonUNC-40 activity. Because of the SDAL process, this inhibition increased nonUNC-40 activity at the anterior surface of the leading edge. At positions 2 and 3, increasing levels of the nonUNC-40 activity at the dorsal and ventral surfaces inhibit UNC-40 activity. The increase in nonUNC-40 activity at the dorsal and ventral surfaces cause a decrease in nonUNC-40 activity at the anterior surface. As a result, the degree to which the direction of nonUNC-40-mediated outgrowth activity fluctuates is greatest at position 3. **(C)** For each position in A, random walk modeling is shown as described in Figure 2E. The response to the extracellular cues progressively increases the degree to which the direction of outgrowth activity fluctuates. By position 3, the degree of fluctuation causes a much lower displacement of membrane mass. The low rate of outward movement causes extension to stall.

We hypothesize that mutations affect the degree to which the direction of nonUNC-40 outgrowth activity fluctuates at position 3 (Figure 15). MIG-15 appears to promote nonUNC-40 SDAL activity, whereas UNC-5 promotes UNC-40 SDAL activity. Because each activity can suppress the other, different domains of nonUNC-40 and UNC-40 SDAL activity can be established along the surface of the leading edge. As the extension moves towards position 3, there is higher nonUNC-40 activity at the more anterior surface and higher UNC-40 activity at the dorsal and ventral surfaces. By suppressing nonUNC 40 activity, the *mig-15* mutation increases, relative to wildtype, the UNC-40 activity at the dorsal and ventral surfaces at position 3. This UNC-40 activity decreases the probability of nonUNC-40 activity at these surfaces and increases the probability of nonUNC-40 activity at the anterior surface. By reducing the degree to which nonUNC-40 outgrowth fluctuates, a greater anterior directional bias is created in the *mig-15* mutants. This results in overextension. The *unc-5* mutation represses the UNC-40 activity at the dorsal and ventral surfaces, increasing the degree to which the direction of the nonUNC-40 outgrowth activity fluctuates. This suppresses the overextension cause by the *mig-15* mutation.

**Figure 15.**
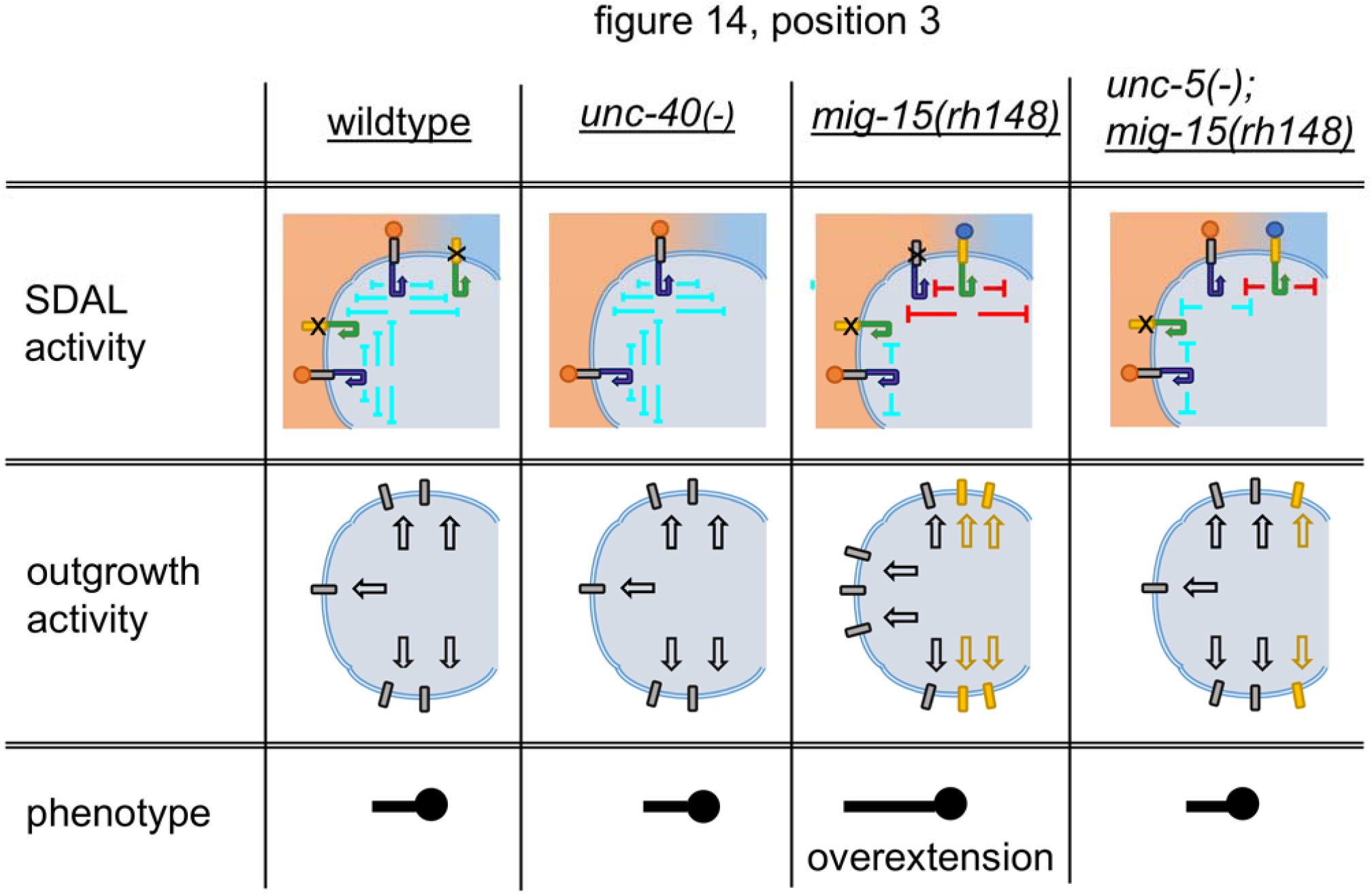
Model for the effects that mutations have on the outgrowth movement of PLM. Table showing the effects of different mutations on the outgrowth of PLM at position 3, Figure 14. PLM outgrowth stalls in wildtype, *unc-40*, and *unc-5;mig-15* mutants at position 3, but overextends in *mig-15* mutants. In this model, the *mig-15* mutation represses the ability of nonUNC-40 SDAL activity to suppress UNC-40 activity at the dorsal and ventral surfaces. Increased UNC-40 SDAL activity suppresses nonUNC-40 SDAL activity at these surfaces. Because of the statistical dependence of the localization process, decreasing nonUNC-40 SDAL activity at the dorsal and ventral surfaces increases nonUNC-40 SDAL activity at the anterior surface. As compared to wildtype, the degree to which the direction of nonUNC-40 outgrowth activity fluctuates is less and, therefore, outward displacement is greater. This allow further anterior outgrowth at position 3. Loss of UNC-5 function in the *mig-15* mutant decreases the ability of UNC-40 SDAL activity to suppresses nonUNC-40 SDAL activity at the dorsal and ventral surfaces, thereby increasing the degree to which the direction of nonUNC-40 outgrowth activity fluctuates. This reduces outward displacement and suppresses the overextension phenotype.

**AVM nerve ring branching and extension phenotype:** Similar to what is proposed for PLM at position 3 in Figure 14, all the surfaces of the leading edge of AVM become exposed to high levels of a cue(s) (Figure 16A, position 1). The degree to which the direction of outgrowth activity fluctuates greatly increases and outward movement stalls. For AVM, this occurs at the nerve ring, which is a source of UNC-6. However, for AVM there are cues at the nerve ring which are arranged perpendicular to one another. We propose that the high level of UNC-6 at all surfaces allows UNC-40 SDAL activity to become more uniformly distributed along all surfaces of the leading edge (Figure 16B) and creates a state where the probabilities of outgrowth in every direction become equal. Both anterior and dorsal outward movement stalls (Figure 16C). This state allows any new cues encountered to effectively create a directional bias (Figure 4). UNC-6 and other cues are arranged along the nerve ring, whereas nonUNC-6 cues are arranged anterior of the nerve ring. As some outgrowth ventures anteriorly and dorsally, these cues stimulate the development of nonUNC-40 and UNC-40 SDAL activity domains (Figure 16B, position 2). Even slight outward movement in the anterior or dorsal directions may reinforce movement in that direction if cues arranged along the axis perpendicular to the direction bias suppress the UNC-40 or nonUNC-40 SDAL activity along the surfaces perpendicular to the directional bias (as depicted in Figure 14B, position 1).

**Figure 16.**
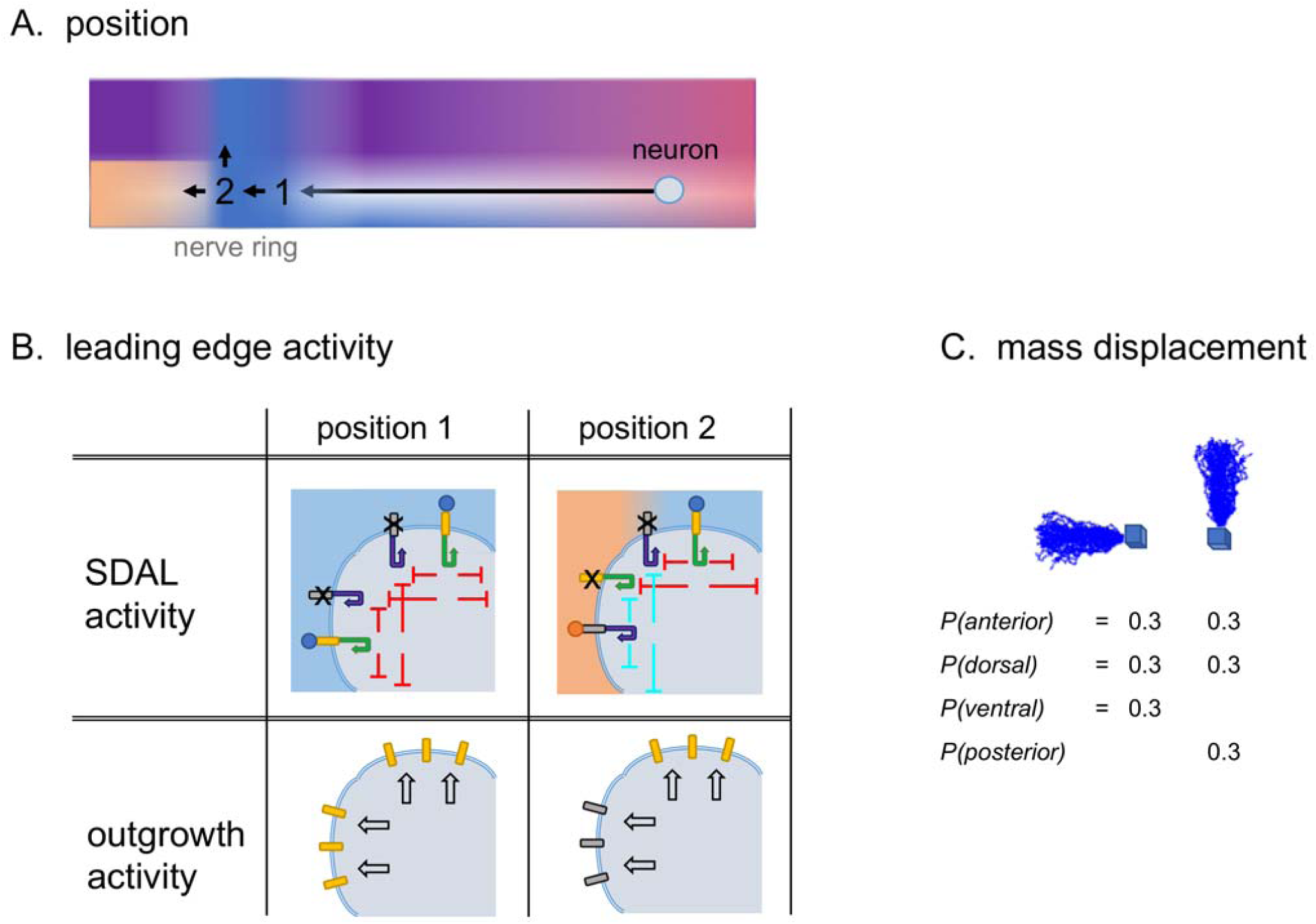
Model for the outgrowth movement of AVM at the nerve ring. Schematic diagrams of the outgrowth of AVM at the nerve ring. The features of the schematic are described in Figure 3. **(A)** At position 1, all surfaces experience high levels of UNC-6. At position 2, the extension encounters new cue(s) at the anterior surface. **(B)** Table showing for each of the positions depicted in A the positive feedback (arrows) and negative feedback (lines) associated with SDAL activity (see Figure 6) and the predicted effect that the SDAL process has on outgrowth activity. At position 1, strong UNC-40 SDAL activity along anterior and dorsal facing surfaces of the leading edge inhibit nonUNC-40 SDAL activity. There is strong UNC-40 mediated outgrowth from all surfaces. At position 2, nonUNC-40 SDAL activity at the anterior surface inhibits UNC-40 SDAL activity, whereas UNC-40 SDAL activity at the dorsal surface inhibits nonUNC-40 SDAL activity. As a result, UNC-40 outgrowth activity is limited to the dorsal surface and nonUNC-40 outgrowth activity is limited to the anterior surface. **(C)** Random walk modeling is shown as described in Figure 2E. Because of the large degree to which the direction of outgrowth fluctuates, outward movement stalls. This allows cues that are arranged dorsal and anterior to the projection to effectively create a bias for outgrowth in the respective direction (see Figure 4). A dorsal directional bias will develop because UNC-40 and other receptors mediates an outgrowth response to UNC-6 and other cues along the nerve ring. An anterior directional bias will develop because nonUNC-40 mediates an outgrowth response to cues along an anterior pathway.

We hypothesize that mutations affect the degree to which the direction of UNC-40 and nonUNC-40 outgrowth activity fluctuates at position 2 (Figure 17). In *unc-40* mutants, the lack of dorsal UNC-40 activity allows the direction of nonUNC-40 outgrowth to fluctuate more. However, the anterior cues increase the probability of anteriorly directed nonUNC-40 outgrowth and, thereby, decrease the probability of dorsally directed activity. This suppresses dorsal extension, while still allowing anterior extension. Loss of MIG-15 activity in the *unc-40* mutant background suppresses the nonUNC-40 SDAL. In comparison to the single *unc-40* mutant, in *unc-40;mig-15* mutants the probability of anterior outgrowth in response to the anterior cues is lower. Consequently, the probability of dorsal nonUNC-40 outgrowth is higher. We speculate that this allows some dorsal extension. In *unc-5* mutants, UNC-40 SDAL activity is reduced, but the activity is still sufficient to allow dorsal extension. Loss of both UNC-5 and MIG-15 function most severely hampers the ability to direct the receptors specifically to one surface. The *unc-5;mig-15* mutants have the most abnormal outgrowth patterns.

**Figure 17.**
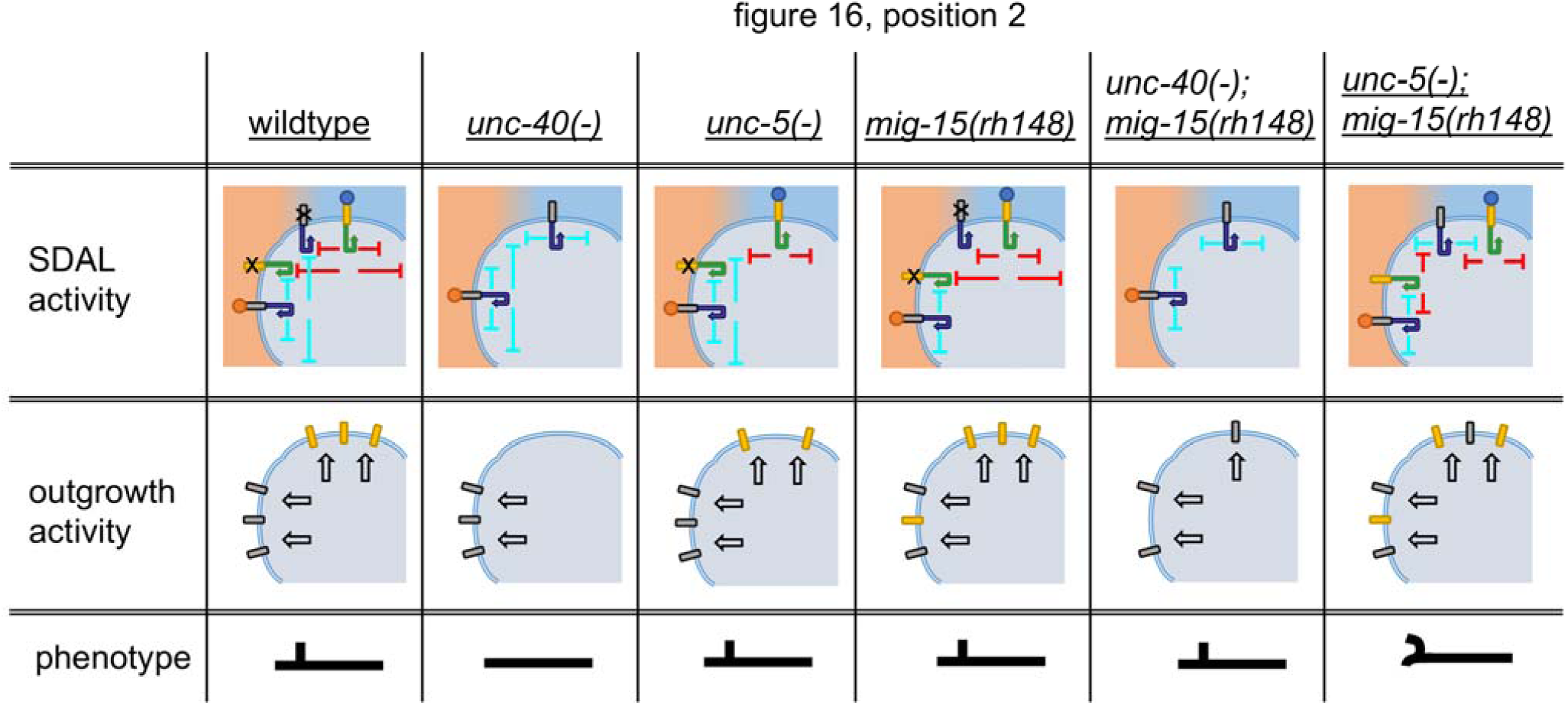
Model for the effects that mutations have on the outgrowth movement of AVM at the nerve ring. Table showing the effects of different mutations on the outgrowth of AVM at position 2, Figure 16. Whereas in wildtype, UNC-40 SDAL activity suppresses nonUNC-40 SDAL activity at the dorsal surface, in *unc-40* mutants there is no suppression and nonUNC-40 activity may occur. However, nonUNC-40 activity is greater at the anterior surface because of the response to anterior cues and this depresses nonUNC-40 activity at the dorsal surface because of the SDAL process. As a result, there is often anterior extension from the nerve ring area but no dorsal extension into the nerve ring. In *unc-5* mutants, UNC-40 SDAL activity is reduced but it is still dorsally oriented. This allows dorsal extension in the mutants. In *mig-15* mutants, nonUNC-40 SDAL activity is repressed. This results in lower nonUNC 40 activity at the anterior surface. However, anterior extension still occur, as does dorsal extension because of UNC-40 activity. Loss of UNC-40 in the *mig-10* background allows extension in both directions. In comparison to the single *unc-40* mutant, the reduced nonUNC-40 SDAL activity at anterior surface in the double *unc-40;mig-15* mutant doesn’t depress dorsal nonUNC-40 outgrowth activity as much, allowing more extension into the nerve ring. Loss of UNC-5 in the *mig-10* background causes the most abnormal outgrowth morphologies, presumably because the repression of both UNC-40 and nonUNC-40 SDAL activities does not allow UNC-40 and nonUNC-40 outgrowth activities to be well sorted to different surfaces.

**HSN extension phenotypes:** We hypothesize that there is high probability for ventrally directed outgrowth from the HSN cell body because of the strong outgrowth-promoting effect of the UNC-40-mediated response to the UNC-6 cue, which is in a higher concentration ventral of the cell body (Figure 18A). We hypothesize that the same process takes place in the HSN neuron as in the PLM neuron, except the movement is towards the UNC-6 source. We depict in Figure 8B that at position 1 there is some nonUNC-40 SDAL activity at the ventral surface of the leading edge. By position 2, higher levels of UNC-6 increase UNC-40 SDAL and suppress ventral nonUNC-40 SDAL activity. At positon 3, UNC-6 is present at high levels along all surfaces and the direction of UNC-40 outgrowth greatly fluctuates. Possibly, the degree to which the direction of UNC-40 and nonUNC-40 outgrowth activity fluctuates is greater at position 1, than at position 2 (Figure 18C). However, at position 3 the fluctuation is greatest.

**Figure 18.**
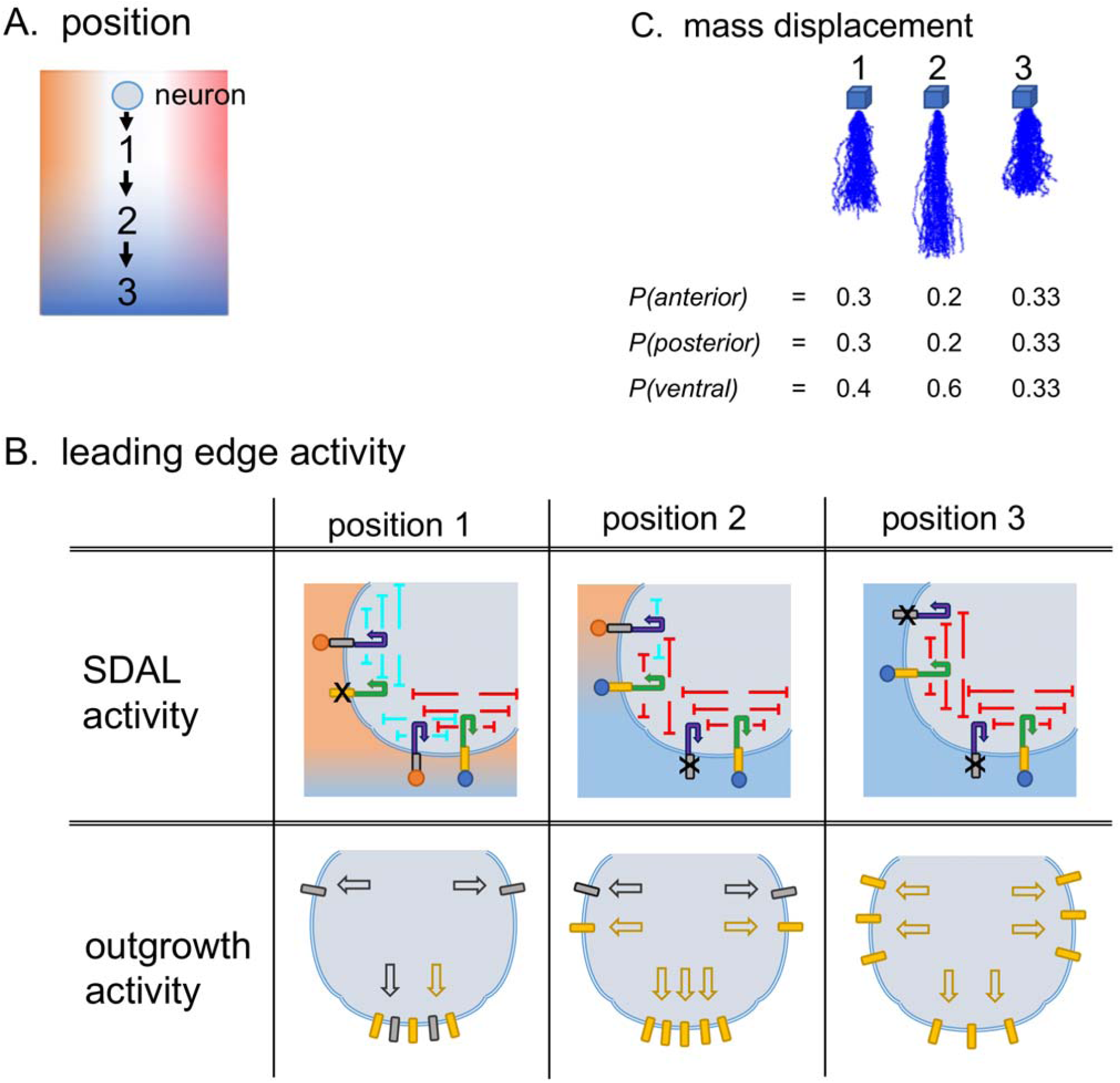
Model for the outgrowth movement of HSN. Schematic diagrams of the ventral outgrowth of HSN. The features of the schematic are described in Figure 3. **(A)** As the leading edge of the extension moves ventrally it encounters higher levels of UNC-6. At position 3, all surfaces experience high levels of UNC-6. Cues anterior and posterior of the pathway maintain an equal probability of outgrowth in both directions. **(B)** Table showing for each of the positions depicted in A the positive feedback (arrows) and negative feedback (lines) associated with SDAL activity (see Figure 6) and the predicted effect that the SDAL process has on outgrowth activity. At position 1, strong UNC-40 SDAL activity along the ventral facing surfaces of the leading edge inhibit nonUNC-40 SDAL activity. Because of the SDAL process, this inhibition increased nonUNC-40 activity at the anterior and posterior (not shown) surfaces of the leading edge. NonUNC-40 SDAL activity along the anterior and posterior surfaces suppress UNC 40 SDAL activity at these surfaces. At positions 2 and 3, increasing levels of the UNC-40 SDAL activity at the anterior and posterior surfaces inhibit nonUNC-40 SDAL activity. The increase in UNC-40 activity at the anterior and posterior surfaces cause a decrease in nonUNC-40 SDAL activity. As a result, the degree to which the direction of UNC-40-mediated outgrowth activity fluctuates is greatest at position 3. **(C)** For each position in A, random walk modeling is shown as described in Figure 2E. At first, the response to the extracellular cues may progressively decreases the degree to which the direction of UNC-40 and nonUNC-40 outgrowth activity fluctuates. However, as UNC-40 SDAL activity predominates at all surfaces, the degree by which the direction of UNC-40 outgrowth fluctuates increases. By position 3, the degree of fluctuation causes a much lower displacement of membrane mass.

We hypothesize that the interplay between UNC-40 and nonUNC-40 SDAL activity allows multiple extensions to develop in the same direction. In fact, UNC-40, UNC-5, MIG-15, and UNC-6 (netrin) activities may function as a type of reaction-diffusion system (TURING 1952; GIERER AND MEINHARDT 1972; MEINHARDT AND GIERER 2000; KONDO AND MIURA 2010; GOEHRING AND GRILL 2013). Along the ventral surface there is a competition between UNC-40 receptors to direct further UNC-40 localization to that site and to inhibit flanking receptors from doing the same. Overtime, the SDAL activity that began at one site predominates, leading to an area of higher outgrowth activity (Figure 19). We speculate that by helping to suppress UNC-40 SDAL activity, nonUNC-40 activity increases the threshold by which the SDAL activity at one site can begin to predominate. The *mig-15* mutation suppresses the nonUNC-40 SDAL activity and decreases the threshold. This may allow more sites along the membrane where UNC-40 SDAL activity can predominate. The sites do not overlap because of the long-range negative feedback that inhibits neighboring UNC-40 activity. The *unc-5* mutation suppresses UNC-40 SDAL activity, both the positive and negative feedback loops. This retards the ability to enhance and localize the process to one area of the surface. This causes greater fluctuation in the direction of outgrowth activity across the entire ventral surface of the neuron. As a result, the rate of initial outgrowth is even less than that which occurs in wildtype and the area of outward growth is more broad.

**Figure 19.**
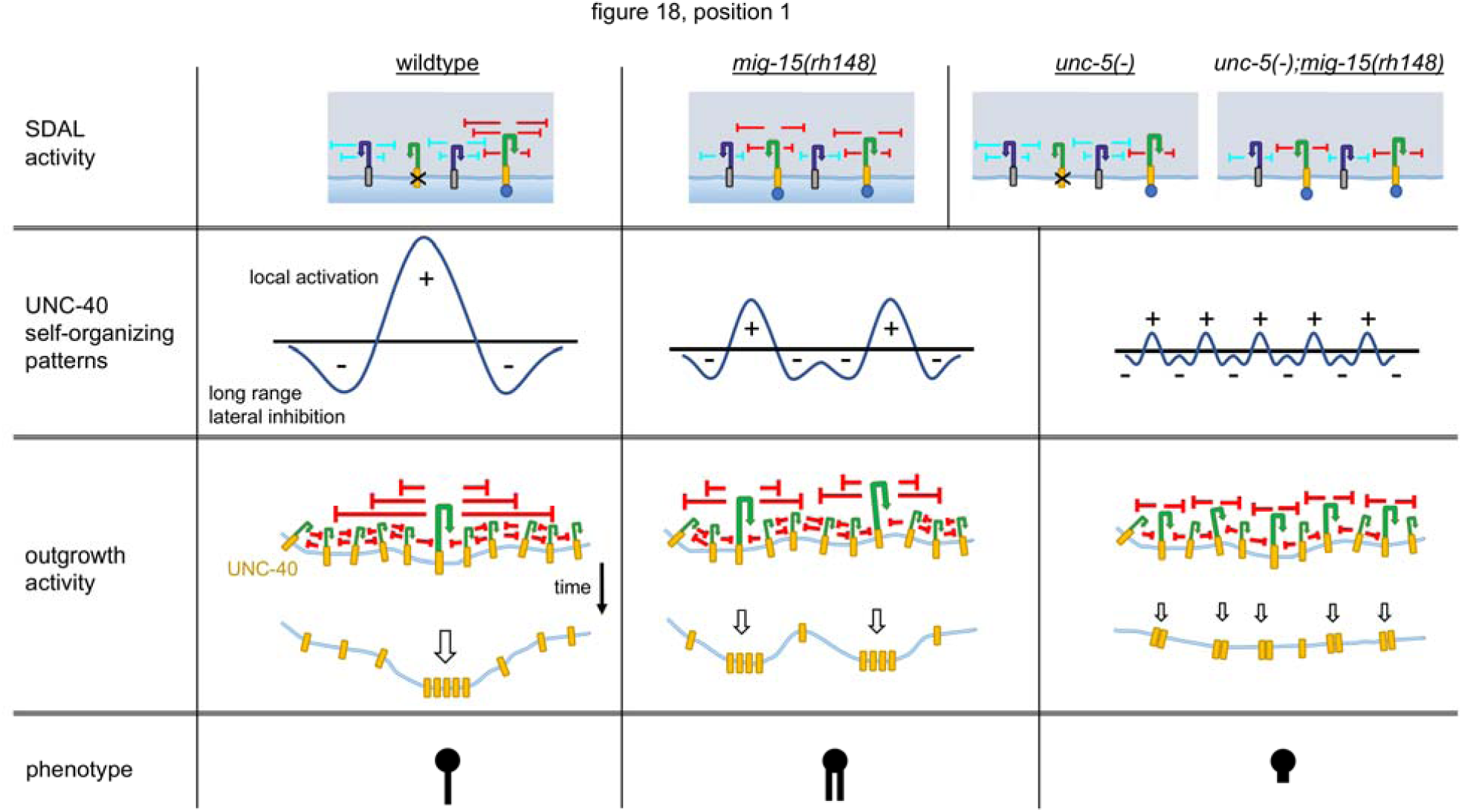
Model for the effects that mutations have on the outgrowth movement of HSN. Table showing the effects of different mutations on the outgrowth of HSN at position 1, Figure 18. Mutations can alter the rate of outgrowth and the number of extensions. In this model, the relative levels of UNC 40 and nonUNC-40 SDAL activity controls this patterning. In wildtype, the ability of UNC-40 SDAL activity to predominate at one site along the membrane is enhanced by nonUNC-40 SDAL activity, which may increase the threshold at which UNC-40 positive feedback becomes effective. Overtime, the area where UNC-40 SDAL predominates causes greater UNC-40 outgrowth activity. As outward movement occurs, higher levels of UNC-6 are encountered. This enhances and localizes the process to that area. In *mig-15* mutants the suppression of nonUNC-40 SDAL activity reduces the threshold at which UNC-40 SDAL activity may predominate at a site. This allows multiple sites along the membrane where UNC-40 SDAL activity may predominate. Multiple extensions may develop as shown in Figure 5. Loss of UNC-5, which suppresses UNC-40 SDAL activity, retards the ability to enhance and localize the process. This causes greater fluctuation in the direction of outgrowth activity across the entire ventral surface of the neuron, which uniformly decreases the displacement of membrane.

### A genetic pathway for UNC-40 asymmetric localization

We present a genetic pathway for the asymmetrical localization of UNC-40 based on the phenotype of robust UNC-40::GFP clustering in HSN. A full understanding of the molecular mechanisms underlying the SDAL process is an important long-term goal. Since we believe that UNC 40::GFP clustering is a readout of that process, constructing genetic pathways for the clustering of UNC-40::GFP is a step toward this goal. We wish to know how UNC-5 mediates signaling within HSN that controls the UNC-40 asymmetric localization process. However, a role for UNC-5 in HSN is paradoxical given the widespread idea that UNC-5 mediates a repulsive response to UNC-6 and that HSN outgrowth is towards the source of UNC-6. All the same, we suggest a cell-autonomous role for UNC-5 in HSN is the most parsimonious model. First, UNC-5 is an UNC-6 receptor that can mediate neuronal responses when in complex with UNC-40 (HONG *et al.* 1999; GEISBRECHT *et al.* 2003; KRUGER *et al.* 2004; FINCI *et al.* 2014). We previously showed that UNC-40 conformational changes regulate HSN asymmetric localization in HSN (XU *et al.* 2009) and we now show that UNC-5 regulates UNC-40 asymmetric localization in HSN. It is therefore plausible that UNC-5 affects UNC-40 conformational changes that regulate UNC-40 asymmetric localization. Second, UNC-5 can alter the number to HSN outgrowths in response to UNC-6 and to the UNC-6ΔC ligand. Directional guidance by UNC-6 and UNC-6ΔC is generally normal in an *unc-5* mutant, suggesting that the ability of UNC-5 to regulate the number of outgrowths is not due to an alteration in the extracellular distribution of its UNC-6 ligand. Further, the UNC-6ΔC ligand and the *mig-15* mutation create the same outgrowth phenotype, which can be suppressed by loss of UNC-5 function, and we have shown that MIG-15 acts cell autonomously in HSN to regulate UNC-40 asymmetric localization (YANG *et al.* 2014). Further, we have shown that the UNC-5-mediated response that regulates UNC-40 asymmetric localization also depends on UNC-53 (NAV2) (KULKARNI *et al.* 2013), a cytoplasmic protein that functions cell-autonomously for cell migration and axon guidance (STRINGHAM *et al.* 2002). Together, these observations strongly suggest that UNC-5 directly regulates signaling within HSN. Third, a role for UNC-5 in the guidance of AVM and PVM axons towards UNC-6 sources has also been suggested. A synergistic interaction between *unc-5* and *egl-20* is observed; in either *unc-5* or *egl-20* mutants the ventral extension of AVM and PVM axons is only slightly impaired, whereas in the double mutants there is a much greater penetrance (LEVY-STRUMPF AND CULOTTI 2014). The expression of an *unc-5* transgene in AVM and PVM can rescue the AVM and PVM axon guidance defects of the *unc-5;egl-20* double mutant (LEVY-STRUMPF AND CULOTTI 2014).

We note that for HSN, transgenic rescue using *unc-5* constructs have not been successful and in wild-type animals UNC-5 expression in HSN has not been reported. As well, expression has not been reported in AVM, PVM, and PLM wild-type neurons. We suspect there may be technical difficulties or that UNC-5 expression might be low in these cells. UNC-5 is detected in PLM in *rpm-1* mutants, which is consistent with evidence that UNC-5 activity is required for PLM overextension in these mutants (LI *et al.* 2008).

To construct genetic pathways, we use the readout of whether UNC-40::GFP is clearly and consistently localized to any side of the HSN neuron in different mutants (Figure 13). A summary of the results is presented (Figure 20A). UNC-6 is required for robust asymmetric UNC-40 localization; in the absence of UNC-6 function UNC-40 remains uniformly distributed along the surface of the plasma membrane. The loss of both UNC-53 and UNC-5 function also results in a uniform distribution, however loss of either one alone does not. This suggests that UNC-53 and UNC-5 pathways act redundantly downstream of UNC-6 (Figure 20B). Moreover, we observe there is robust asymmetric UNC-40 localization when there is a loss of UNC-6 activity in addition to the loss of UNC-53 and UNC-5. This suggests a third pathway that is suppressed by UNC-6 when UNC-53 and UNC-5 activity are missing. Loss of both UNC-5 and UNC-6 does not allow UNC-40 localization, whereas loss of both UNC-53 and UNC-6 does, therefore UNC-53, rather than UNC-5, acts with UNC-6 to suppress the third pathway.

**Figure 20.**
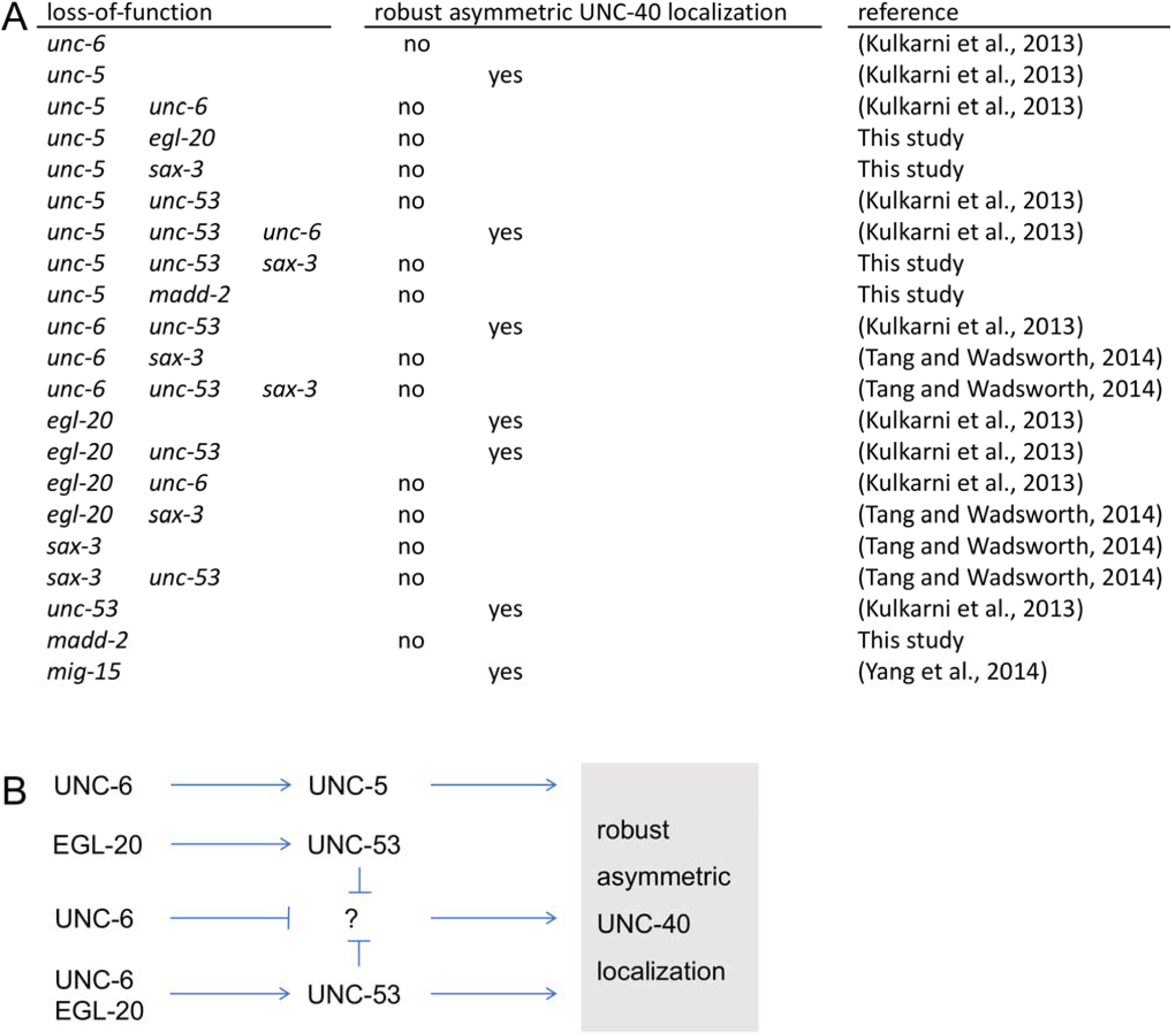
Genetic pathways for self-organizing UNC-40 asymmetric localization. **(A)** Table summarizing the results of experiments previously reported and described in Figure 13 of this paper. **(B)** The genetic data support a model whereby the UNC-6 and EGL-20 extracellular cues regulate at least three pathways leading to robust asymmetric UNC-40 localization. Robust asymmetric UNC-40 localization refers to the ability to observe UNC-40::GFP clustering at the surface of the neuron. Arrows represent activation; bars represent repression. See text for the logic used to construct the pathways.

UNC-40 becomes localized when EGL-20 activity is lost. As well, UNC-40 becomes localized when both EGL-20 and UNC-53 activities are lost. This is consistent with UNC-6 promoting UNC-40 localization via the UNC-5 pathway. Loss of EGL-20 and UNC-5 prevents UNC-40 localization. In these animals, the UNC-5 pathway is absent and UNC-6 is present to block the third pathway, therefore the UNC-53 pathway that leads to UNC-40 localization must require EGL-20, as well as UNC-6.

Loss of UNC-6 activity or loss of both UNC-6 and EGL-20 activity prevents localization, whereas loss of only EGL-20 does not. To explain this, we propose that when UNC-6 is lost, the third pathway, which would otherwise be activated by the loss of UNC-6, remains suppressed because EGL-20 activity promotes suppression via UNC-53 activity. This suppression also explains why loss of UNC-6 and UNC-5 activity does not cause localization.

The genetic pathways are consistent with the models proposed in Figures 1 and 6. In the models, positive feedback loops amplify the polarized responses to extracellular cues, whereas negative feedback limits the responses and confines the positive feedback to the sites of interaction. We hypothesize that the UNC-5 and UNC-53 genetic pathways shown at the top and bottom of Figure 20B correspond to the positive feedback loops depicted in Figures 1 and 6 by the arrows. The “?” genetic pathway corresponds to UNC-40 asymmetric localization and outgrowth activity in the absence of UNC-6. The UNC-53 genetic pathways in Figure 20B that block the “?” pathway corresponds to the negative feedback loops (lines) in Figures 1 and 6 which prevent UNC-40 asymmetric localization and outgrowth in the absence of UNC-6. Loss of both UNC-6 and EGL-20 prevents robust asymmetric UNC-40 localization because both UNC-6- and EGL-20-mediated positive feedback loops are disrupted. A positive feedback loop may be necessary to establish a negative feedback loop. Therefore, the “?” pathway is not active when both UNC-6 and EGL-20 are absent.

Importantly, this genetic analysis indicates that netrin (UNC-6) and wnt (EGL-20) signaling are integrated to regulate self-organizing UNC-40 asymmetric localization. An implication of this result is that the extracellular concentrations of UNC-6 and EGL-20 could control the activation or inhibition of UNC-40-mediated outgrowth. This could be important for generating patterns of outgrowth when neurons move to new locations within the animal. The picture is complicated by the evidence that both UNC-6 and EGL-20 affect the SDAL of both UNC-40 mediated and nonUNC-40-mediated outgrowth activity. It is possible that overlapping sets of extracellular cues and their receptors are involved in setting the probability of outgrowth for each activity. SAX-3 and MADD-2 are required for UNC-40::GFP localization, but also affect nonUNC-40-mediated outgrowth. The *egl-20;sax-3* and *unc-40;sax-3* double mutations have the greatest effect on restricting the extent of outgrowth movement in any direction (Figures 6 and 7). Moreover, the number of HSN neurites is reduced in *unc-5* mutants, whereas the number in *unc-5;sax-3* double mutants appears normal (Figure 7B). Understanding the interdependence of these outgrowth activities could provide a better understanding of how extracellular cues affect the patterns of outgrowth *in vivo*.

## Acknowledgments

We thank Caenorhabditis Genetics Center, J. Culotti, and C. Bargmann for strains; we thank Martha Soto and members of the Soto laboratory for support and helpful discussions; we thank Martha Soto, Peter Yurchenco, Bhumi Patel and Leely Rezvani for comments on the manuscript. We also thank the editors and reviewers of the journal for their insightful comments and suggestions. This work was supported by grants NS033156 and NS061805 from the National Institutes of Health, National Institute of Neurological Disorders and Stroke and grant 07-3060 SCR-E-0 from the New Jersey Commission on Spinal Cord to WGW. This work was also supported by grant DFHS13PPCO28 from the New Jersey Commission on Cancer Research to AM.

## References

Adler, C. E., R. D. Fetter and C. I. Bargmann, 2006 UNC-6/Netrin induces neuronal asymmetry and defines the site of axon formation. Nat Neurosci 9: 511–518.

Aiello, G. L., 2016 Neuro-Percolation as a Superposition of Random-Walks, pp. in European Symposium on Artificial Neural Networks, Computational Intelligence and Machine Learning. i6doc.com Bruges, Belgium.

Alexander, M., K. K. Chan, A. B. Byrne, G. Selman, T. Lee et al., 2009 An UNC-40 pathway directs postsynaptic membrane extension in Caenorhabditis elegans. Development 136: 911–922.

Alexander, M., G. Selman, A. Seetharaman, K. K. Chan, S. A. D’Souza et al., 2010 MADD-2, a homolog of the Opitz syndrome protein MID1, regulates guidance to the midline through UNC-40 in Caenorhabditis elegans. Developmental Cell 18: 961–972.

Arrieumerlou, C., and T. Meyer, 2005 A local coupling model and compass parameter for eukaryotic chemotaxis. Dev Cell 8: 215–227.

Asakura, T., K. Ogura and Y. Goshima, 2007 UNC-6 expression by the vulval precursor cells of Caenorhabditis elegans is required for the complex axon guidance of the HSN neurons. Dev Biol 304: 800–810.

Baier, H., and F. Bonhoeffer, 1992 Axon guidance by gradients of a target-derived component. Science 255: 472–475.

Bourne, H. R., and O. Weiner, 2002 A chemical compass. Nature 419: 21.

Buettner, H. M., R. N. Pittman and J. K. Ivins, 1994 A model of neurite extension across regions of nonpermissive substrate: simulations based on experimental measurement of growth cone motility and filopodial dynamics. Dev Biol 163: 407–422.

Chan, S. S., H. Zheng, M. W. Su, R. Wilk, M. T. Killeen et al., 1996 UNC-40, a C. elegans homolog of DCC (Deleted in Colorectal Cancer), is required in motile cells responding to UNC-6 netrin cues. Cell 87: 187–195.

Chang, C., C. E. Adler, M. Krause, S. G. Clark, F. B. Gertler et al., 2006 MIG-10/Lamellipodin and AGE-1/PI3K Promote Axon Guidance and Outgrowth in Response to Slit and Netrin. Curr Biol 16: 854–862.

Chapman, J. O., H. Li and E. A. Lundquist, 2008 The MIG-15 NIK kinase acts cell-autonomously in neuroblast polarization and migration in C. elegans. Dev Biol 324: 245–257.

Colavita, A., and J. G. Culotti, 1998 Suppressors of ectopic UNC-5 growth cone steering identify eight genes involved in axon guidance in Caenorhabditis elegans. Dev Biol 194: 72–85.

Finci, L. I., N. Krüger, X. Sun, J. Zhang, M. Chegkazi et al., 2014 The crystal structure of netrin-1 in complex with DCC reveals the bifunctionality of netrin-1 as a guidance cue. Neuron 83: 839–849.

Fraser, A. G., R. S. Kamath, P. Zipperlen, M. Martinez-Campos, M. Sohrmann et al., 2000 Functional genomic analysis of C. elegans chromosome I by systematic RNA interference. Nature 408: 325–330.

Geisbrecht, B. V., K. A. Dowd, R. W. Barfield, P. A. Longo and D. J. Leahy, 2003 Netrin binds discrete subdomains of DCC and UNC5 and mediates interactions between DCC and heparin. J Biol Chem 278: 32561–32568.

Gierer, A., and H. Meinhardt, 1972 A theory of biological pattern formation. Kybernetik 12: 30–39.

Goehring, N. W., and S. W. Grill, 2013 Cell polarity: mechanochemical patterning. Trends Cell Biol 23: 72–80.

Gopal, A. A., B. Rappaz, V. Rouger, I. B. Martyn, P. D. Dahlberg et al., 2016 Netrin-1-Regulated Distribution of UNC5B and DCC in Live Cells Revealed by TICCS. Biophys J 110: 623–634.

Graziano, B. R., and O. D. Weiner, 2014 Self-organization of protrusions and polarity during eukaryotic chemotaxis. Curr Opin Cell Biol 30: 60–67.

Hagedorn, E. J., J. W. Ziel, M. A. Morrissey, L. M. Linden, Z. Wang et al., 2013 The netrin receptor DCC focuses invadopodia-driven basement membrane transmigration in vivo. J Cell Biol 201: 903–913.

Hao, J. C., C. E. Adler, L. Mebane, F. B. Gertler, C. I. Bargmann et al., 2010 The tripartite motif protein MADD-2 functions with the receptor UNC-40 (DCC) in Netrin-mediated axon attraction and branching. Dev Cell 18: 950–960.

Hao, J. C., T. W. Yu, K. Fujisawa, J. G. Culotti, K. Gengyo-Ando et al., 2001 C. elegans Slit Acts in Midline, Dorsal-Ventral, and Anterior-Posterior Guidance via the SAX-3/Robo Receptor. Neuron 32: 25–38.

Harterink, M., D. H. Kim, T. C. Middelkoop, T. D. Doan, A. van Oudenaarden et al., 2011 Neuroblast migration along the anteroposterior axis of C. elegans is controlled by opposing gradients of Wnts and a secreted Frizzled-related protein. Development 138: 2915–2924.

Hedgecock, E. M., J. G. Culotti and D. H. Hall, 1990 The unc-5, unc-6, and unc-40 genes guide circumferential migrations of pioneer axons and mesodermal cells on the epidermis in C. elegans. Neuron 4: 61–85.

Hong, K., L. Hinck, M. Nishiyama, M. M. Poo, M. Tessier-Lavigne et al., 1999 A ligand-gated association between cytoplasmic domains of UNC5 and DCC family receptors converts netrin-induced growth cone attraction to repulsion. Cell 97: 927–941.

Katz, M. J., E. B. George and L. J. Gilbert, 1984 Axonal elongation as a stochastic walk. Cell Motil 4: 351–370.

Keleman, K., and B. J. Dickson, 2001 Short- and long-range repulsion by the Drosophila Unc5 netrin receptor. Neuron 32: 605–617.

Killeen, M., J. Tong, A. Krizus, R. Steven, I. Scott et al., 2002 UNC-5 function requires phosphorylation of cytoplasmic tyrosine 482, but its UNC-40-independent functions also require a region between the ZU-5 and death domains. Dev Biol 251: 348–366.

Kondo, S., and T. Miura, 2010 Reaction-diffusion model as a framework for understanding biological pattern formation. Science 329: 1616–1620.

Kruger, R. P., J. Lee, W. Li and K. L. Guan, 2004 Mapping netrin receptor binding reveals domains of Unc5 regulating its tyrosine phosphorylation. J Neurosci 24: 10826–10834.

Kulkarni, G., Z. Xu, A. M. Mohamed, H. Li, X. Tang et al., 2013 Experimental evidence for UNC-6 (netrin) axon guidance by stochastic fluctuations of intracellular UNC-40 (DCC) outgrowth activity. Biol Open 2: 1300–1312.

Lai Wing Sun, K., J. P. Correia and T. E. Kennedy, 2011 Netrins: versatile extracellular cues with diverse functions. Development 138: 2153–2169.

Leung-Hagesteijn, C., A. M. Spence, B. D. Stern, Y. Zhou, M. W. Su et al., 1992 UNC-5, a transmembrane protein with immunoglobulin and thrombospondin type 1 domains, guides cell and pioneer axon migrations in C. elegans. Cell 71: 289–299.

Levy-Strumpf, N., and J. G. Culotti, 2014 Netrins and Wnts function redundantly to regulate antero-posterior and dorso-ventral guidance in C. elegans. PLoS Genet 10: e1004381.

Li, H., G. Kulkarni and W. G. Wadsworth, 2008 RPM-1, a Caenorhabditis elegans protein that functions in presynaptic differentiation, negatively regulates axon outgrowth by controlling SAX-3/robo and UNC-5/UNC5 activity. J Neurosci 28: 3595–3603.

Lim, Y. S., S. Mallapur, G. Kao, X. C. Ren and W. G. Wadsworth, 1999 Netrin UNC-6 and the regulation of branching and extension of motoneuron axons from the ventral nerve cord of Caenorhabditis elegans. J Neurosci 19: 7048–7056.

MacNeil, L. T., W. R. Hardy, T. Pawson, J. L. Wrana and J. G. Culotti, 2009 UNC-129 regulates the balance between UNC-40 dependent and independent UNC-5 signaling pathways. Nat Neurosci 12: 150–155.

Maes, T., A. Barcelo and C. Buesa, 2002 Neuron navigator: a human gene family with homology to unc-53, a cell guidance gene from Caenorhabditis elegans. Genomics 80: 21–30.

Manser, J., C. Roonprapunt and B. Margolis, 1997 C. elegans cell migration gene mig-10 shares similarities with a family of SH2 domain proteins and acts cell nonautonomously in excretory canal development. Dev Biol 184: 150–164.

Manser, J., and W. B. Wood, 1990 Mutations affecting embryonic cell migrations in Caenorhabditis elegans. Dev Genet 11: 49–64.

Maskery, S. M., H. M. Buettner and T. Shinbrot, 2004 Growth cone pathfinding: a competition between deterministic and stochastic events. BMC Neurosci 5: 22.

McShea, M. A., K. L. Schmidt, M. L. Dubuke, C. E. Baldiga, M. E. Sullender et al., 2013 Abelson interactor-1 (ABI-1) interacts with MRL adaptor protein MIG-10 and is required in guided cell migrations and process outgrowth in C. elegans. Dev Biol 373: 1–13.

Meinhardt, H., and A. Gierer, 2000 Pattern formation by local self-activation and lateral inhibition. Bioessays 22: 753–760.

Merz, D. C., H. Zheng, M. T. Killeen, A. Krizus and J. G. Culotti, 2001 Multiple signaling mechanisms of the unc-6/netrin receptors unc-5 and unc-40/dcc in vivo. Genetics 158: 1071–1080.

Moore, S. W., M. Tessier-Lavigne and T. E. Kennedy, 2007 Netrins and their receptors. Adv Exp Med Biol 621: 17–31.

Morikawa, R. K., T. Kanamori, K. Yasunaga and K. Emoto, 2011 Different levels of the Tripartite motif protein, Anomalies in sensory axon patterning (Asap), regulate distinct axonal projections of Drosophila sensory neurons. Proc Natl Acad Sci U S A 108: 19389–19394.

Mortimer, D., T. Fothergill, Z. Pujic, L. J. Richards and G. J. Goodhill, 2008 Growth cone chemotaxis. Trends Neurosci 31: 90–98.

Mortimer, D., Z. Pujic, T. Vaughan, A. W. Thompson, J. Feldner et al., 2010 Axon guidance by growth-rate modulation. Proc Natl Acad Sci U S A 107: 5202–5207.

Nguyen, H., P. Dayan and G. J. Goodhill, 2014 The influence of receptor positioning on chemotactic information. J Theor Biol 360: 95–101.

Nguyen, H., P. Dayan and G. J. Goodhill, 2015 How receptor diffusion influences gradient sensing. Journal of The Royal Society Interface 12.

Norris, A. D., and E. A. Lundquist, 2011 UNC-6/netrin and its receptors UNC-5 and UNC-40/DCC modulate growth cone protrusion in vivo in C. elegans. Development.

Odde, D. J., and H. M. Buettner, 1995 Time series characterization of simulated microtubule dynamics in the nerve growth cone. Ann Biomed Eng 23: 268–286.

Pan, C. L., J. E. Howell, S. G. Clark, M. Hilliard, S. Cordes et al., 2006 Multiple Wnts and frizzled receptors regulate anteriorly directed cell and growth cone migrations in Caenorhabditis elegans. Dev Cell 10: 367–377.

Poinat, P., A. De Arcangelis, S. Sookhareea, X. Zhu, E. M. Hedgecock et al., 2002 A conserved interaction between beta1 integrin/PAT-3 and Nck-interacting kinase/MIG-15 that mediates commissural axon navigation in C. elegans. Curr Biol 12: 622–631.

Quinn, C. C., D. S. Pfeil, E. Chen, E. L. Stovall, M. V. Harden et al., 2006 UNC-6/netrin and SLT-1/slit guidance cues orient axon outgrowth mediated by MIG-10/RIAM/lamellipodin. Curr Biol 16: 845–853.

Quinn, C. C., D. S. Pfeil and W. G. Wadsworth, 2008 CED-10/Rac1 mediates axon guidance by regulating the asymmetric distribution of MIG-10/lamellipodin. Curr Biol 18: 808–813.

Ren, X. C., S. Kim, E. Fox, E. M. Hedgecock and W. G. Wadsworth, 1999 Role of netrin UNC-6 in patterning the longitudinal nerves of Caenorhabditis elegans. J Neurobiol 39: 107–118.

Rosoff, W. J., J. S. Urbach, M. A. Esrick, R. G. McAllister, L. J. Richards et al., 2004 A new chemotaxis assay shows the extreme sensitivity of axons to molecular gradients. Nat Neurosci 7: 678–682.

Sawa, H., and H. C. Korswagen, 2013 Wnt signaling in C. elegans. WormBook: 1–30.

Shakir, M. A., J. S. Gill and E. A. Lundquist, 2006 Interactions of UNC-34 Enabled with Rac GTPases and the NIK kinase MIG-15 in Caenorhabditis elegans axon pathfinding and neuronal migration. Genetics 172: 893–913.

Sloan, T. F., M. A. Qasaimeh, D. Juncker, P. T. Yam and F. Charron, 2015 Integration of shallow gradients of Shh and Netrin-1 guides commissural axons. PLoS Biol 13: e1002119.

Song, S., Q. Ge, J. Wang, H. Chen, S. Tang et al., 2011 TRIM-9 functions in the UNC-6/UNC-40 pathway to regulate ventral guidance. Journal of genetics and genomics = Yi chuan xue bao 38: 1–11.

Stringham, E., N. Pujol, J. Vandekerckhove and T. Bogaert, 2002 unc-53 controls longitudinal migration in C. elegans. Development 129: 3367–3379.

Stringham, E. G., and K. L. Schmidt, 2009 Navigating the cell: UNC-53 and the navigators, a family of cytoskeletal regulators with multiple roles in cell migration, outgrowth and trafficking. Cell adhesion & migration 3: 342–346.

Tang, X., and W. G. Wadsworth, 2014 SAX-3 (Robo) and UNC-40 (DCC) Regulate a Directional Bias for Axon Guidance in Response to Multiple Extracellular Cues. PLoS One 9: e110031.

Tessier-Lavigne, M., and C. S. Goodman, 1996 The molecular biology of axon guidance. Science 274: 1123–1133.

Teulière, J., C. Gally, G. Garriga, M. Labouesse and E. Georges-Labouesse, 2011 MIG-15 and ERM-1 promote growth cone directional migration in parallel to UNC-116 and WVE-1. Development 138: 4475–4485.

Turing, A. M., 1952 The Chemical Basis of Morphogenesis.

Wadsworth, W. G., H. Bhatt and E. M. Hedgecock, 1996 Neuroglia and pioneer neurons express UNC-6 to provide global and local netrin cues for guiding migrations in C. elegans. Neuron 16: 35–46.

Wadsworth, W. G., and E. M. Hedgecock, 1996 Hierarchical guidance cues in the developing nervous system of C. elegans. Bioessays 18: 355–362.

Wang, F. S., C. W. Liu, T. J. Diefenbach and D. G. Jay, 2003 Modeling the role of myosin 1c in neuronal growth cone turning. Biophys J 85: 3319–3328.

Wang, Z., Y. Hou, X. Guo, M. van der Voet, M. Boxem et al., 2013 The EBAX-type Cullin-RING E3 ligase and Hsp90 guard the protein quality of the SAX-3/Robo receptor in developing neurons. Neuron 79: 903–916.

Wang, Z., L. M. Linden, K. M. Naegeli, J. W. Ziel, Q. Chi et al., 2014 UNC-6 (netrin) stabilizes oscillatory clustering of the UNC-40 (DCC) receptor to orient polarity. J Cell Biol 206: 619–633.

Whangbo, J., and C. Kenyon, 1999 A Wnt signaling system that specifies two patterns of cell migration in C. elegans. Molecular cell 4: 851–858.

Tang, X., and W.G. Wadsworth, 2014 SAX-3 (Robo) and UNC-40 (DCC) Regulate a Directional Bias for Axon Guidance in Response to Multiple Extracellular Cues.

Xu, Y., H. Taru, Y. Jin and C. C. Quinn, 2015 SYD-1C, UNC-40 (DCC) and SAX-3 (Robo) function interdependently to promote axon guidance by regulating the MIG-2 GTPase. PLoS Genet 11: e1005185.

Xu, Z., H. Li and W. G. Wadsworth, 2009 The roles of multiple UNC-40 (DCC) receptor-mediated signals in determining neuronal asymmetry induced by the UNC-6 (netrin) ligand. Genetics 183: 941–949.

Yang, Y., W. S. Lee, X. Tang and W. G. Wadsworth, 2014 Extracellular Matrix Regulates UNC-6 (Netrin) Axon Guidance by Controlling the Direction of Intracellular UNC-40 (DCC) Outgrowth Activity. PLoS One 9: e97258.

Yoshimura, S., J. I. Murray, Y. Lu, R. H. Waterston and S. Shaham, 2008 mls-2 and vab-3 Control glia development, hlh-17/Olig expression and glia-dependent neurite extension in C. elegans. Development 135: 2263–2275.

Zallen, J. A., B. A. Yi and C. I. Bargmann, 1998 The conserved immunoglobulin superfamily member SAX-3/Robo directs multiple aspects of axon guidance in C. elegans. Cell 92: 217–227.

Ziel, J. W., E. J. Hagedorn, A. Audhya and D. R. Sherwood, 2009 UNC-6 (netrin) orients the invasive membrane of the anchor cell in C. elegans. Nat Cell Biol 11: 183–189.

